# FASTAptameR 2.0: A Web Tool for Combinatorial Sequence Selections

**DOI:** 10.1101/2022.04.27.489774

**Authors:** Skyler T. Kramer, Paige R. Gruenke, Khalid K. Alam, Dong Xu, Donald H. Burke

## Abstract

Combinatorial selections are powerful strategies for identifying biopolymers with specific biological, biomedical, or chemical characteristics. Unfortunately, most available software tools for high-throughput sequencing analysis have high entrance barriers for many users because they require extensive programming expertise. FASTAptameR 2.0 is an R-based reimplementation of FASTAptamer designed to minimize this barrier while maintaining the ability to answer complex sequence-level and population-level questions. This opensource toolkit features a user-friendly web tool, interactive graphics, up to 100x faster clustering, an expanded module set, and an extensive user guide. FASTAptameR 2.0 accepts diverse input polymer types and can be applied to any sequence-encoded selection.

## Background

Combinatorial selections are powerful strategies for identifying biopolymers with specific characteristics such as target specificity or affinity, catalytic properties, or biological function. The strength and adaptability of this approach were recognized with the 2018 Nobel Prize in Chemistry for Francis Arnold, George Smith, and Gregory Winter [1]. While these biopolymers are generally composed of nucleotides or amino acids, the molecular alphabets can be extended or modified to include non-canonical amino acids [2] and chemically modified nucleotides such as AEGIS [3], Hachimoji [4], and others [5]. Selection strategies for nucleic acids have been applied to aptamers [6,7], (deoxy)ribozymes [8–10], synthetic genetic polymers (XNAs) [11,12], and other combinatorial chemistries. Selection strategies for peptides and proteins can be accomplished by selecting for bioactivity in cells or whole organisms [13] or by display on phage particles [14], ribosomes [15], mRNA [16], whole bacteria [17], and other platforms. The genes that encode the evolving proteins can be translated from nucleic acid libraries according to the standard genetic code or to natural or artificial genetic codes [18]. DNA sequence libraries have even been used as barcodes to track lipid nanoparticle formulations [19–21] and combinatorial chemical synthesis [22,23]. In short, any platform that links polymer sequence (genotype) with a selectable or screenable property (phenotype) can be adapted to combinatorial selections.

Under optimal circumstances, the evolutionary dynamics of populations undergoing selection reflect the relative fitness of each species, with high-fitness sequences typically enriching during selection and low-fitness sequences depleting. Thus, common analytic tasks of any combinatorial selection include counting the number of occurrences for each sequence [24,25], calculating enrichment of sequences between two or more rounds [25–27], filtering sequences based on the number of reads present in one or multiple rounds [28], clustering related sequences [25,29–31], and in some cases analyzing predicted structure motifs [29,32–36]. High-throughput sequencing (HTS) provides large volumes of data for these analyses and can yield high-resolution insights. Many specialized bioinformatics toolkits have been developed to enable this analytical workflow [37], and several of these tools include graphical user interfaces to visualize HTS data during the analysis [36,38,39]. However, some of these toolkits require significant computational resources or coding expertise that together constitute barriers to entry for the average molecular biologist. Our lab previously developed and released the FASTAptamer toolkit [25] to address the primary, sequence-level needs in the field, such as those outlined above. FASTAptamer is an open-source toolkit consisting of five Perl scripts that can be used to count, normalize, and rank reads in a FASTQ or FASTA file; compare populations for sequence distribution; cluster related sequences; calculate fold-enrichment; and search for sequence motifs [25]. Since its publication, the FASTAptamer toolkit has been used and cited extensively for diverse types of molecular and biological selections on populations of functional nucleic acids and protein/peptides (see **Additional file 1**), thereby demonstrating its ability to address many of the first-level bioinformatics needs of the field.

Although FASTAptamer can analyze sequences from many types of biomolecules, its original application was targeted to aptamers, which are structured nucleic acids capable of binding to a molecular target, usually with high specificity and affinity. Aptamers are generated through an iterative, *in vitro* selection process termed SELEX (Systematic Evolution of Ligands by EXponential enrichment) [6,7]. After a determined number of selection rounds, the sequences of the enriched aptamers have traditionally been obtained by cloning the aptamer libraries into a plasmid and sequencing each clone one at a time.

With HTS, millions of sequence reads from multiple rounds of selection can be determined, and this information can be used to identify aptamer candidates for further characterization [40–47]. HTS investigation of *in vitro* selection pools has revealed the distribution and relative frequencies of individual sequences and groups of related sequences as the populations evolve through the course of the experiment [48,49]. Such data can inform on the success of the selection [45,50], aptamer-target interactions [51–54], the mutation and fitness landscape [29,44,55], structure-function relations, biological constraints, and more.

While the initial release of FASTAptamer is generally user-friendly, it also has some limitations. First, as the FASTAptamer modules are Perl scripts, they must be run using a command line, which creates a modest barrier for practitioners of combinatorial selections who are unfamiliar or uncomfortable with a command-line interface. Second, depending on the parameters used, the clustering module is time-consuming and computationally intensive [31]. Third, while the output data from FASTAptamer can be downloaded for offline visualization, it does not allow for visualization of results within the platform, which can constrain data exploration. To address these limitations, we describe here the development of FASTAptameR 2.0, an R-based reimplementation of FASTAptamer. This program improves upon the original version while keeping the features that made FASTAptamer an accessible, easy-to-use toolkit for the analysis of HTS datasets. Like FASTAptamer, FASTAptameR 2.0 does not need external dependencies (especially when used through the web tool) and is easy to install and launch. FASTAptameR 2.0 is portable across multiple platforms, open source, and comes with a detailed user guide that includes screenshots of the user interface and sample output tables and graphs for each module (see **Additional file 2).** Further, the generalizable outputs can be used as downstream inputs to this program or any other bioinformatics program that supports FASTA files. It has a userfriendly interface that can be accessed online at https://fastaptamer2.missouri.edu/ or in a downloadable form as a Docker image from Docker Hub (https://hub.docker.com/repository/docker/skylerkramer/fastaptamer2), and the code can be accessed from GitHub at https://github.com/SkylerKramer/FASTAptameR-2.0 [56]. Additional improvements in FASTAptameR 2.0 include a faster clustering algorithm with speeds nearly 100X faster than FASTAptamer in some cases (*e.g.,* for larger, more complex libraries) and an expanded set of interconnected modules (shown in **Fig. 1)** that can be used to interactively analyze and visualize HTS data from new perspectives with custom, user-defined pipelines. Collectively, these improvements make exploration of HTS data from combinatorial selections significantly more accessible.

**Figure 1.**
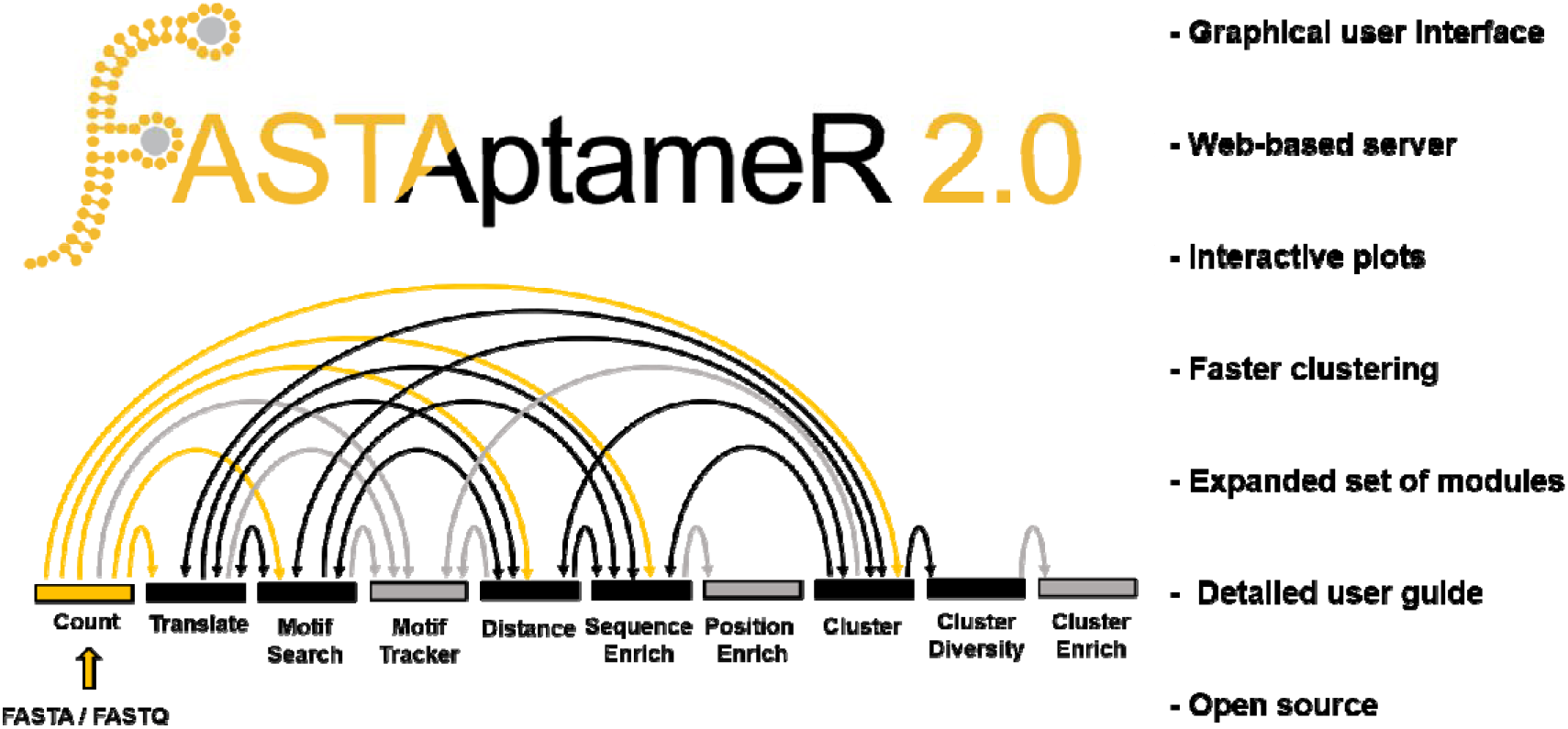
General overview, module connectivity, and major new features of FASTAptameR 2.0. All workflows start with the Count module (gold node). Outputs from mid-stage modules (black nodes) can be inputs for other modules. Outputs from end-stage modules (gray nodes) are not used elsewhere within FASTAptameR 2.0. Output tables and calculations from most modules can be downloaded as FASTA, CSV, or both.

## Results

### Count module

The first step in analyzing sequence data from combinatorial selections is nearly always to determine the read count (abundance) of each unique sequence. This information can indicate whether the population is relatively diverse with little convergence or has converged on one or a few dominant sequences. Either of these scenarios is immediately evident when analyzing the population with the *Count* module, which, as in FASTAptamer, is the entry point into FASTAptameR 2.0.

This module serves two main purposes. First, it condenses the original file size by returning a FASTA file with a single entry for each unique sequence. Second, it provides summary statistics for each unique sequence in the input population as three key metrics: abundance, rank by abundance, and reads per million (RPM), which is the read count divided by the population size in millions. It then incorporates these statistics into the sequence identifier for each entry. For example, a sequence with an identifier of “>4-94978-43966.9” is the fourth most abundant sequence in its population, has 94,978 identical reads, and is present at a frequency of 43,966.9 RPM. The distribution of those statistical values across a given population provides the first insights into the degree to which that population has converged, which can be seen by visualizing the relationship between rank and abundance **(Fig. 2a,** generated with the 70HRT14 population from the original FASTAptamer publication [51,57–59]; see **Materials and Methods).** A slowly decreasing function suggests that the population has not converged onto a small set of sequences, whereas a steeply decreasing line suggests convergence.

**Figure 2.**
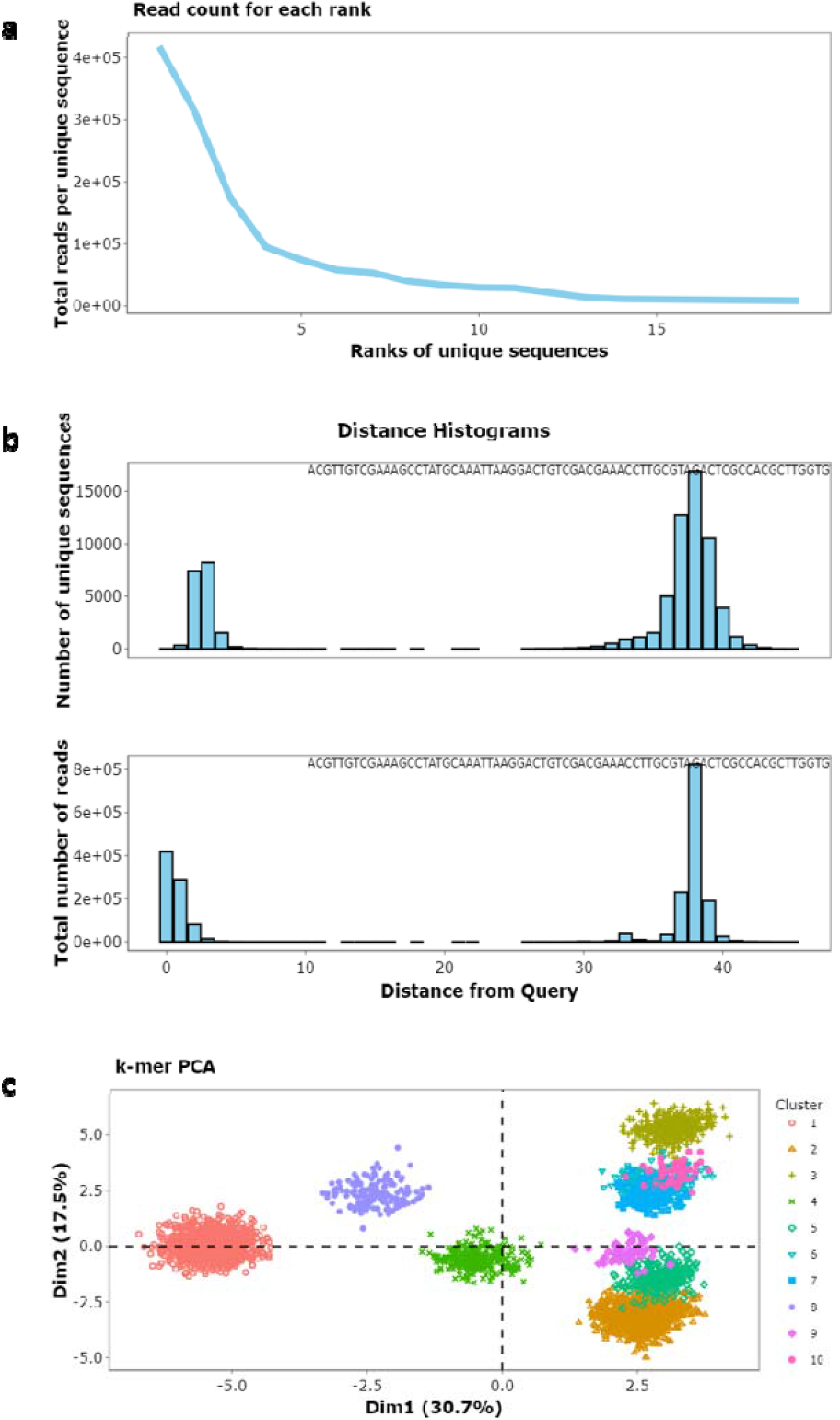
Example plots in a Cluster module workflow, using the 70HRT14 population as an example. a) This line plot of the relationship between sequence rank and abundance (from the Count module) suggests that the population is dominated by a few sequences (convergence) due to its relatively steep slope and the magnitude of the y-axis. b) These histograms of LEDs suggest that many sequences in this population are similar to its most abundant sequence and that the region of sequence space surrounding the most abundant sequence is well-sampled, which can indicate biochemical significance. For the top plot, each unique sequence is equally weighted, whereas each unique sequence in the bottom plot is weighted by abundance. c) A 3-mer matrix was generated from clustered sequences and visualized as a PCA plot where colors correspond to clusters.

### Distance module

The *Distance* module is a new feature of FASTAptameR 2.0 that computes the Levenshtein edit distance (LED) between a query sequence and every other sequence in the population. An in-house precursor of this module was previously used in an *in vivo* selection [60]. Distance analysis can be especially useful when monitoring the accumulation of point mutants, evaluating the effectiveness of a mutagenesis protocol, or monitoring diversity near the beginning of a selection from sequences that densely sample local sequence space (*e.g.,* via mutagenic PCR or doped resynthesis). The distribution of these LEDs can then be visualized as a histogram of distances (**Fig. 2b**) to provide additional perspectives on overall sequence relatedness within the population. For output libraries, an isolated cluster will be seen close to zero distance when the population consists predominantly of sequences that are closely related to the query (such as when the query is part of a cluster of sequences that have come to dominate the population) or after the accumulation of new mutations during the course of the selection (drift or divergent evolution from of the founder sequences). A second peak will appear at large distances from the query when the remaining sequences are evolutionarily unrelated to the query, such as when many different founding members of a random sequence population are independently selected **(Fig. 2b).** To illustrate, this analysis was applied to the 70HRT14 population, using the most abundant sequence as the query. Plotting this distribution *by equally weighting each unique sequence* (top plot of **Fig. 2b**) reveals that many of these sequences are similar to the query, that the region of sequence space surrounding this query is well-sampled (which can indicate biochemical significance), that most species are within three mutations relative to the query, and that nearly all sequences related to the query are within an LED of 7. This visualization provides guidance in setting the maximum LED value to use in the *Cluster* module (next section). In contrast, plotting the distribution after *weighting the data by sequence abundance* (bottom plot of **Fig. 2b)** shows that variants within one or two mutations of the query are far more abundant than those with higher-order mutations.

### Cluster algorithm and validation

The *Cluster* module groups closely related sequences into ‘clusters’, thereby setting the stage for computing local fitness landscapes and further simplifying downstream analysis. The clustering algorithms for FASTAptamer and FASTAptameR 2.0 are both iterative, greedy processes that start by considering the most abundant, unclustered sequence as a cluster seed. Given that the *Count* module sorts the population by abundance, the first sequence in that output because the first cluster seed here. All unique, unclustered sequences within a predetermined LED are added to this cluster. After considering all unique sequences in the population, the next most abundant, unclustered sequence becomes the seed of the second cluster, and all unclustered sequences within the LED are added to that cluster. These steps are iterated until a predetermined stop condition is met (see below).

Clustering can be computationally intense and slow, a problem that has been observed for FASTAptamer and other platforms [31]. FASTAptameR 2.0 significantly reduces the clustering runtime by changing the underlying data structure for the computations. The original implementation stores clustered sequences in arrays (a static data structure), whereas the FASTAptameR 2.0 implementation uses linked lists (a dynamic data structure). The list structure more efficiently handles memory requirements, which grow with the size and complexity of the population. FASTAptameR 2.0 offers additional means of reducing clustering time, such as by allowing the user to filter out sequences with abundance less than a user-defined threshold or to set a maximum number of clusters for the module to generate.

The FASTAptameR 2.0 clustering algorithm was benchmarked against FASTAptamer by comparing runtimes on an Ubuntu subsystem (v18.04) on a desktop computer with 16 GB RAM and an Intel I7 processor. All 72,921 unique sequences from the 70HRT14 population were used to generate the top thirty clusters with a maximum LED of seven. While the original implementation finished in 35 minutes 27 seconds, the FASTAptameR 2.0 implementation finished in 24.6 seconds - roughly 86 times faster. When clustering times were compared for a number of other scenarios, FASTAptameR 2.0 was always significantly faster than FASTAptamer, and the magnitude of this difference grew with the size and complexity of the population being analyzed. Therefore, the FASTAptameR 2.0 clustering algorithm is strictly better than the FASTAptamer one.

Outputs from the *Cluster* module can be visualized with the *Cluster Diversity* module, which is another new addition to FASTAptameR 2.0. This module uses all unique sequences within each of the user-defined clusters to create a k-mer matrix. This matrix is subsequently visualized as a two-dimensional PCA plot **(Fig. 2c).** The k-mer plot for the top ten clusters of the 70HRT14 population shows most clusters in well-defined regions with separation from most or all of the other clusters. The separations among the clusters reflect their origins from independent, unrelated founder sequences present in the initial population, while the spread of each cluster reflects the sampling of local sequence space around each founder sequence, resulting from the accumulation of functional point mutations through neutral neutral drift and purifying selection.

Sample plots from a cluster-specific workflow are shown in **Fig. 2.** The panels show evidence of a convergent population **(Fig. 2a**), quantify the distance from a query sequence to the rest of the population and suggest LED values required for clustering **(Fig. 2b**), and provide a visualization of the separation between and diversity within clusters **(Fig. 2c).**

### Individual and population-level tracking

Another new feature of FASTAptameR 2.0 is the ability to track individual motifs, sequences, or clusters across multiple rounds of a selection experiment. The *Motif Tracker* module tracks query motifs or sequences, and the *Enrich* module tracks how every sequence changes between populations, while the *Cluster Enrich* module tracks whole clusters to allow the user to monitor the collective behavior of the cluster as a whole, analogous to the collective evolutionary dynamics of a viral quasispecies. These three modules additionally calculate how families or species enrich, which can be indicative of how they performed in the selection experiment. An in-house precursor of the *Motif Tracker* module was previously used in a selection for 2’-modified RNA aptamers with affinity for HIV-1 RT [61]. In the case when two clustered files are supplied to the *Enrich* module, the enrichment values of individual sequences can be grouped by clustering and visualized as a box plot **(Fig. 3a).** Cluster 2 of **Fig. 3a** is enriched relative to the other clusters, suggesting that this set of sequences has a motif important for target binding (specifically the family 1 pseudoknot, or F1Pk, in this case). For each cluster in the box plot, individual points that are well above or well below the median represent species that are enriching or depleting relative to the cluster as a whole, both of which can be highly informative - species with strongly advantageous variations may be emerging as future dominant species for that cluster, while species with strongly disadvantageous variations can illuminate critical portions of the biomolecule, as illustrated by the next module.

**Figure 3.**
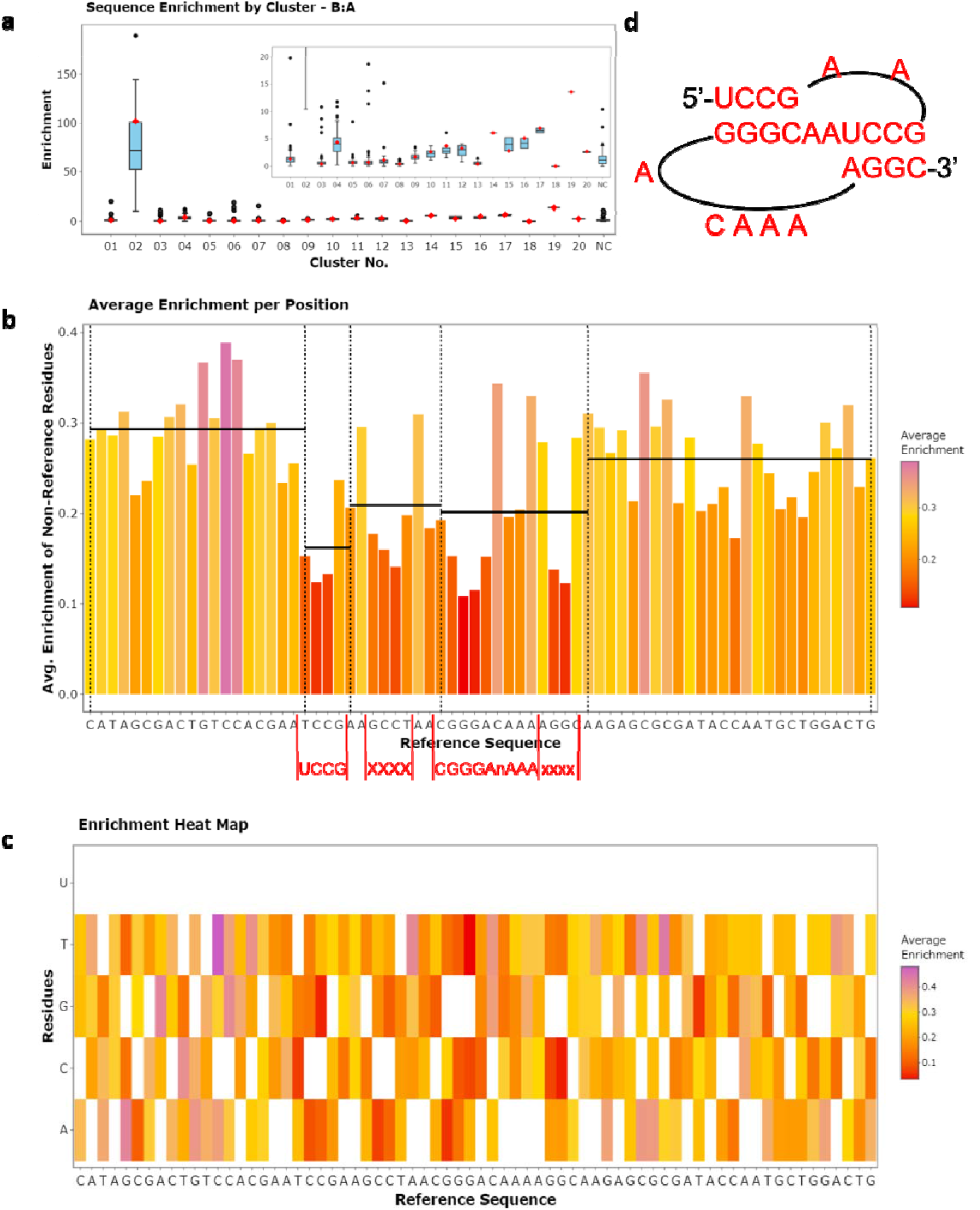
Cluster and position enrichment plots. a) The cluster box plot showing how clustered sequences in 70HRT14 enrich in 70HRT15. Cluster 2 of 70HRT14, for example, is highly enriched in 70HRT15 due to the presence of the F1Pk, which is implicated in target binding to HIV-1 RT. The 25th and 75th quartiles are respectively represented by the bottom and top of each box. The line in the middle of the box represents the median. Whiskers are at most 1.5 * IQR (interquartile range), and any points beyond that are shown as outliers. The red marker indicates where the seed sequence of the cluster falls. b) The x-axis of the bar plot also shows the user-defined reference sequence, and the y-axis shows the average enrichment of each non-reference residue at each position. The red text below this panel shows the portion of the query sequence that matches the linear F1Pk motif. Black, horizontal lines show the left-inclusive average enrichment score of each user-defined region. The regions corresponding to the F1Pk motif have the lowest regional average of enrichment scores, indicating the importance of this motif for this selection experiment. c) The x-axis of the heat map shows the user-defined reference sequence, and they-axis shows all possible residues at each position. Colors depict the average enrichment of each possible non-reference residue. d) The experimentally determined secondary structure of the F1Pk motif.

### Position enrichment

For a set of closely related sequences, mutations in some positions contribute directly to enrichment or depletion, while mutations at some other positions have little consequence. Uncovering these relationships can be enormously valuable for delineating the contributions of those positions to macromolecular functions. The *Position Enrichment* module is a new feature of FASTAptameR 2.0 that calculates the average enrichment or depletion at each position for all sequences that do not match the corresponding user-defined reference residue at that position. This calculation is visualized as a bar plot **(Fig. 3b).** Relatively short bars indicate functional conservation at that position, such that deviations from the reference residue identity contribute to depletion. Exceptionally tall bars may indicate positive selection for improved function relative to the reference sequence. As a result, highly conserved sections are immediately visible as regions with low bars because mutations in these positions contribute to depletion. The module calculates local averages across user-defined intervals and displays them as horizontal lines across those intervals, making the conserved and non-conserved regions especially evident from visual inspection **(Fig. 3b).** *Position Enrichment* further resolves enrichment and depletion patterns for each of the available substitute residues (*e.g.,* 3 alternative nucleotides or 19 alternative amino acids when using standard alphabets) and displays the resulting patterns as a heat map **(Fig. 3c).** As in the *Translate* module, nonstandard nucleotides and amino acids can be analyzed with the *Position Enrichment* module.

To generate the plots in **Fig. 3,** the 70HRT14 and 70HRT15 populations were counted and clustered following the workflow of **Fig. 2,** although in this case the *Count* module was also used to omit any sequences that were not exactly 70 nucleotides long. The *Enrich* module created the box plot from the full set of clustered sequences in both populations. The *Enrich* module then calculated enrichment scores scores for the first cluster from 70HRT15, which carries the F1Pk motif, and for the corresponding cluster from the preceding round of selection (second cluster from 70HRT14) [58,62]. The *Position Enrichment* module used the output from the *Enrich* module as input, and the most abundant sequence in 70HRT15 was used as the reference sequence. The segments within the 70-nucleotide random region that contain the pseudoknot at the functional core of the aptamer are shown in **Fig. 3d.**

### Expanded sequence support

While populations of any sequence type (*e.g.,* nucleotides or amino acids) could be fed into the original release of FASTAptamer, it did not allow for sequence translations. The *Translate* module of FASTAptameR 2.0 translates nucleotide sequences to amino acids according to either the standard genetic code or any of fifteen alternative genetic codes such as those used by vertebrate mitochondria, mycoplasma, and other organisms. This module also supports complete customization of the genetic code used for translation to support nonstandard nucleotide input and/or nonstandard amino acid output, both of which are useful for applications in synthetic biology. Thus, FASTAptameR 2.0 explicitly supports all linear biopolymers of diverse biological origins.

## Discussion

The integration of HTS with combinatorial selections experiments has created many opportunities and challenges for bioinformatics analyses. Though many tools exist to aid these analyses [24,25,28,30,31,34–36,38,39], usually they are not designed for users without a strong computational background. As such, the user may need to devote significant time and effort to tasks such as software installation and dependency handling before they can even learn to properly use the tool, constituting a serious barrier to data exploration.

FASTAptameR 2.0 was designed with these users in mind and according to best practices in the field of bioinformatics [63–66]. Like its predecessor FASTAptamer, FASTAptameR 2.0 is a powerful open-source toolkit to analyze combinatorial selection populations and is accompanied by an extensive user guide. The program simplifies data analysis by minimally requiring a web browser and internet access. Further, the outputs are designed to be modular so that this platform can be easily integrated into existing workflows or used to develop custom ones. Module inputs and outputs are standard file types (*e.g.,* FASTQ/A and CSV), and the UI can be accessed from any browser operating with any operating system. It can also be run locally through any system with a functional Docker installation.

Modules in this platform can be used for a wide range of functions on subsets of individual populations or across many populations. *FASTAptameR-Count,* the starting point of the platform, counts and ranks unique sequences. *FASTAptameR-Translate* translates nucleotide sequences according to standard, nonstandard, or user-defined genetic codes. *FASTAptameR-Motif_Search* and *FASTAptameR-Motif_Tracker* modules identify occurrences of motifs and track them across populations over time. *FASTAptameR-Distance* computes the LED between all sequences and a query sequence. *FASTAptameR-Enrich* and *FASTAptameR-Position_Enrich* assess sequence trajectories across populations and provide insights into which residues contribute to the enrichment scores. The three linked modules of *FASTAptameR-Cluster, FASTAptameR-Cluster_Diversity,* and *FASTAptameR-Cluster_Enrich* cluster sequences, provide cluster-level metadata, and assess how they change across populations.

FASTAptameR 2.0 features substantial improvements relative to its predecessor. By increasing user accessibility, improving the original modules, and providing additional tools for data analysis, FASTAptameR 2.0 further lowers the technical barrier for analysis and exploration of HTS datasets and allows the user to gain more insights from their combinatorial selection experiments.

## Materials and methods

### Data description

Data from the original FASTAptamer publication [51,57,58] were used to build and test FASTAptameR 2.0. In brief, these data are two populations of RNA aptamers selected to target HIV-1 reverse transcriptase after 14 and 15 rounds of a SELEX experiment (designated 70HRT14 and 70HRT15, respectively). These populations were trimmed via cutadapt [67] and filtered for high-quality reads via the FASTX-Toolkit [68]. These FASTQ files are available in [59].

### Implementation

FASTAptameR 2.0 is written in the R programming language [69] and made interactive with the Shiny package [70]. The platform uses ggplot2 [71] to build plots and plotly [72] to make them interactive. The entire program (*i.e.*, code, dependencies, and supporting files) is wrapped into a Docker image and deployed on a web server at the University of Missouri - Columbia. The web server has been tested in Google Chrome, FireFox, and Safari. Beta testers at five institutions confirmed platform independence and the absence of external dependencies.

## Declarations

### Availability of data and materials

All code and supporting files are available at https://github.com/SkylerKramer/FASTAptameR-2.0 [56]. The Docker image is available at https://hub.docker.com/repository/docker/skylerkramer/fastaptamer2. Finally, the web-accessible version of FASTAptameR 2.0 is available at https://fastaptamer2.missouri.edu/. All data analyzed in this manuscript are available in [59].

FASTAptameR 2.0 is distributed under a GNU General Public License version 3.0.

### Authors’ contributions

STK developed the frontend and backend, prepared the tool for deployment, interacted with beta testers, and wrote the manuscript and user guide with input from PRG, KKA, DX, and DHB. DHB supervised the project, recruited and interacted with beta testers, conceived the project with PRG, and edited the manuscript. The authors read and approved the final manuscript.

## Acknowledgements

Feedback from beta testing was provided by Austin Prater, Marc Johnson, Carolyn Robinson, and Laurie Agosto from the University of Missouri - Columbia; Uli Müller from the University of California - San Diego; Andrej Luptak from the University of California - Irvine; and Elisa Biondi from the Foundation for Applied Molecular Evolution. The authors thank Rebecca Burke-Agüero for critical discussions and technical assistance early in the development of a faster clustering algorithm and Matt Stanley for network support during launch. The authors also thank members of the Burke and Xu research teams and the NASA Interdisciplinary Consortium for Astrobiology Research ‘Bringing RNA to Life’ (NASA grant 80NSSC21K0596) for feedback, discussion, and advice throughout the project.

## Supplementary information

*Additional file 1: Table of FASTAptamer use cases.* Word document that contains a table summarizing many use cases of the original FASTAptamer publication.

*Additional file 2: FASTAptameR 2.0 user guide.* PDF document that contains the user guide for FASTAptameR 2.0.

