## Additional file 1 for "FASTAptameR 2.0: A Web Tool for Combinatorial Sequence Selections"

Additional text for Introduction:

Since its publication, the FASTAptamer toolkit has been used and cited extensively for diverse types of molecular and biological selections on populations of functional nucleic acids and proteins/peptides (Table S1), thereby demonstrating its ability to address many of the first-level bioinformatics needs of the field.

**Table S1: FASTAptamer Use Cases**

|  | Type of Selection | Description of Study | FASTAptamer Modules Used | Reference |
| --- | --- | --- | --- | --- |
| Nucleic Acids | DNA Aptamers | Cell-SELEX to CD44E/s | Count | [1] |
|  |  | Selection to A549 cells for aptamers that internalize | Count, Cluster | [2] |
|  |  | Selection to multidrug-resistant carcinoma | Did not specify | [3] |
|  |  | Selection to *A. muciniphila* gut bacteria | Count, Compare, Enrich | [4] |
|  |  | Selection to carbapenem resistant *P. aeruginosa* | Count, Cluster, Enrich | [5] |
|  |  | Identification of aptamer to human monocytes and macrophages | Count, Enrich | [6] |
|  |  | Ligand-guided SELEX to TCR-CD3ε | Count, Enrich | [7] |
|  |  | Ligand-guided SELEX with AEGIS components to TCR-CDε | Count, Enrich | [8] |
|  |  | Selection to clear cell renal cell carcinoma RCC-MF cell line | Count, Enrich | [9] |
|  |  | Cell-SELEX to T-cell marker CD8 | Count, Enrich | [10] |
|  |  | Selection to CTLA-4 | Did not specify | [11] |
|  |  | Selection to bovine spermatozoa | Count, Cluster | [12] |
|  |  | Selection to human TNFα | Count, Cluster | [13] |
|  |  | Selection to LAG3 | Count, Cluster | [14] |
|  |  | Selection to human PD-L1 | Did not specify | [15] |
|  |  | Selection to the murine extracellular domain of PD-1 | Did not specify | [16] |
|  |  | Selection to human adenovirus | Count, Enrich | [17] |
|  |  | Selection to N-terminal domain of SARS-CoV-2 spike protein | Did not specify | [18] |
|  |  | Selection to Newcastle avian virus for detection applications | Count, Cluster | [19] |
|  |  | Selection to HMG1 of *Plasmodium falciparum* | Count, Enrich | [20] |
|  |  | Selection to Galectin-1 | Did not specify | [21] |
|  |  | Selection to kanamycin for development of structure-switching aptamer-based biosensor | Count, Cluster, Compare, Enrich | [22] |
|  |  | Selection to diverse synthetic cathinones | Did not specify | [23] |
|  |  | Selection to serotonin as part of single-walled carbon nanotube (SWCNT) sensor | Did not specify | [24] |
|  | RNA Aptamers | Selection (2ʹFY, 2ʹdA RNA) to G-protein coupled receptors (GPCRs) | Count, Enrich | [25] |
|  |  | Re-selection to diverse strains of HIV-1 RT | Count, Cluster, Compare, Enrich, Search | [26] |
|  |  | 2ʹ-modified RNA re-selection to HIV-1 RT | Count, Cluster, Enrich, Search | [27] |
|  |  | Mannose-modified RNA selection to lectin concanavalin A | Count, Enrich | [28] |
|  |  | Selections to thrombin and TGFβ1 | Count, Cluster | [29] |
|  |  | 2ʹFY selection to MRP1 used to create MRP1-CD28 bispecific aptamer | Did not specify | [30] |
|  |  | 2ʹFY selection to TIM3 | Count, Cluster | [31] |
|  | (Deoxy)Ribozymes | Selection for RNA ligase activity using libraries with different-sized random region | Count, Compare, Cluster | [32] |
|  |  | Tenofovir-transferase ribozyme selection | Count, Cluster, Enrich | [33] |
|  |  | Endonuclease deoxyribozyme selection | Count, Cluster, Enrich | [34] |
|  |  | Self-cleaving ribozyme selection in presence of a mineral surface | Count, Cluster, Compare | [35] |
|  |  | Self-cleaving ribozyme selection in diverse chemical environments | Count, Cluster, Compare | [36] |
|  | Nucleobase editing | Development of base-editing assay coupled with HTS | Count, Enrich | [37] |
|  |  | Used assay above to compare system using tethered APOBEC3B (A3B) to the Cas9n/gDNA complex and MagnEdit system when an A3B-interacting protein is tethered to the Cas9n/gDNA complex and recruits APOBEC3B to targeted edit sites | Count, Enrich | [38] |
|  | Miscellaneous | Determine small RNA loading specificity onto *Drosophila* Argonaute proteins | Count, Enrich | [39] |
| Peptides and Proteins | Phage Display | Selection to traumatic brain injury biomarkers | Count, Enrich | [40] |
|  |  | Selection of phagebodies that was started by mutagenizing the host-range-determining-region (HRDR) of phage tail fibers | Count | [41] |
|  | Biological activity *in vivo* | Screening of HIV-1 Vif mutant library | Count, Enrich | [42] |
|  |  | Screening of MLV Env mutant library | Count, Enrich | [43] |
|  | mRNA/cDNA Display | Proof-of-principle selection to anti-FLAG M2 antibody identified consensus FLAG epitope motif | Count | [44] |
