## Additional file 2 for "FASTAptameR 2.0: A Web Tool for Combinatorial Sequence Selections"

### FASTAptameR 2.0 - User Interface Tutorial

2022/February/11

**4 Version history** **46**

**References** **47**

### 1 Introduction

FASTAptameR 2.0 expands the bioinformatics pipeline that was first released as FASTAptamer (Alam KK 2015). Like its predecessor, FASTAptameR 2.0 is an open-source toolkit designed to quickly and easily analyze populations of sequences resulting from combinatorial selections. This updated version is an R-based platform that features a graphical user interface (UI), interactive graphics, more modules, and a faster implementation of the original clustering algorithm.

This user guide walks through installation of the software and operations for each of the modules, and it highlights what options are available to analyze data through the UI.

The key features of the ten modules of FASTAptameR 2.0 are outlined below in **Section 1.1**. **Fig. 1** gives an overview of how these modules connect to one another. Additionally, a summary of input and output file types is given in **Table 1**. Please note that each module requires the user to upload a file or, in the case of **FASTAptameR-Count**, optionally provide a GitHub link to the data. At present, none of these data will be saved on the server to be passed between modules.

A feature of FASTAptameR 2.0 is that many function inputs/outputs are simply FASTA files, so FASTAptameR 2.0 can be easily integrated into most analytical pipelines. Note that *counted* FASTA files are the minimum input for most modules (*e.g.*, FASTAptameR-Translate needs *at least* a counted FASTA from FASTAptameR-Count but could also accept a searched or clustered FASTA file).

Importantly, FASTAptameR 2.0 does not provide any functions that are easily addressed by other software (*e.g.*, merging paired-end reads, trimming constant regions, predicting structures, *etc.*). Rather, the focus of this application is to provide flexible downstream analyses that are especially applicable to, and useful for, the combinatorial selections field.

#### 1.1 Overview

- **FASTAptameR-Count**
  - This module is the entry point into FASTAptameR 2.0
  - Input: preprocessed FASTQ/A
  - Workflow:
    1. count the occurrence of each unique sequence (`Reads`)
    2. sort by counts in descending order (`Rank`)
    3. normalize counts to reads per million (`RPM`)
  - Interactive plotting:
    1. line plot of reads for each unique sequence sorted by rank
    2. histograms of sequence lengths - one for the unique sequences and one for all reads
    3. sequence abundance bar plot
  - Output: FASTA or CSV (Note: FASTA output from this module is referred to as 'counted FASTA' files throughout this tutorial)‘
- **FASTAptameR-Translate**
  - Input: counted FASTA from FASTAptameR-Count
  - Workflow: translate D/RNA sequences to amino acid sequences
  - Interactive plotting:
    1. line plot of reads for each unique sequence sorted by rank
    2. histograms of sequence lengths - one for the unique sequences and one for all reads
  - Output: FASTA or CSV
- **FASTAptameR-Motif_Search**
  - Input: counted FASTA from FASTAptameR-Count and user-defined, comma-separated query patterns

  - Workflow: search for user-defined query patterns in sequences
  - Output: FASTA or CSV

- **FASTAptameR-Motif_Tracker**
  - Input: at least two counted FASTA files from FASTAptameR-Count and query list (either motifs or full sequences)
  - Workflow: track how user-defined motifs or sequences from the query list change across populations
  - Interactive plotting: line plot of each query's RPM across the populations
  - Output: CSV to summarize each query + CSV of enrichment values

- **FASTAptameR-Distance**
  - Input: counted FASTA from FASTAptameR-Count and query sequence
  - Workflow: compute the Levenshtein edit distance (LED) between the query sequence and all other provided sequences
  - Interactive plotting: histograms of edit distances - one for the unique sequences and one for all reads
  - Output: FASTA or CSV

- **FASTAptameR-Enrich**
  - Input: at least two counted FASTA files from FASTAptameR-Count or two FASTA files from FASTAptameR-Cluster (below), which will be referred to as "clustered FASTA" files
  - Workflow: calculate how each sequence enriches across populations
  - Interactive plotting:
    1. bar plot of sequence persistence
    2. histogram(s) of `log2(Enrichment)`
    3. scatter plot(s) of RPM
    4. Ratio average (RA) plot, displaying average log-RPM (`A`) and log ratio (`R`)
    5. box plot(s) of sequence enrichment per cluster; only available if clustered FASTA files from FASTAptameR-Cluster are provided
  - Output: CSV

- **FASTAptameR-Position_Enrich**
  - Input: enrichment CSV from FASTAptameR-Enrich and reference sequence
  - Workflow: for each position of the reference sequence, compute the average enrichment of non-reference residues in the data
  - Interactive plotting:
    1. bar plot of average enrichment per position of reference sequence
    2. heat map of average enrichment per position of reference sequence grouped by residues
  - Output: No direct file output (only interactive plots)

- **FASTAptameR-Cluster**
  - Input: counted FASTA from FASTAptameR-Count
  - Workflow:
    1. filter out low-read sequences based on user-defined input
    2. treat the most abundant, non-clustered sequence as cluster seed
    3. add all sequences within a user-defined LED of the seed to the cluster
    4. Repeat Steps 2 and 3 until all sequences are clustered or a maximum number of clusters are created
  - Output: FASTA or CSV (Note: FASTA output from this module is referred to as 'clustered FASTA' files throughout this tutorial)'

- **FASTAptameR-Cluster_Diversity**

  - Input: clustered FASTA from FASTAptameR-Cluster
  - Workflow: provide metadata for each cluster
  - Interactive plotting:
    1. metaplots for count of unique sequences, count of total reads, and average LED per cluster
    2. k-mer PCA plot, colored by cluster identity
  - Output: CSV

- **FASTAptameR-Cluster_Enrich**
  - Input: at least two CSVs from FASTAptameR-Cluster_Diversity
  - Workflow: calculate how each cluster enriches across populations
  - Interactive plotting: line plot of each the total RPM per cluster for each seed per population
  - Output: CSV to summarize each cluster + CSV of enrichment values

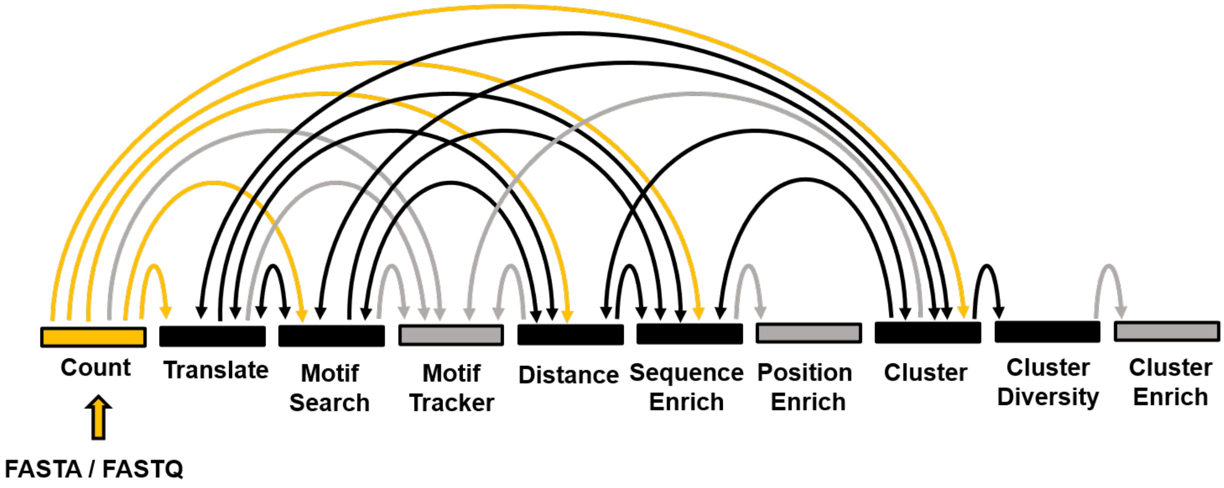

Figure 1: Arc Diagram of Module Connections. All workflows must start with the Count module (gold). Black modules (nodes) have outputs that can be used elsewhere, whereas outputs from gray modules (nodes) are not able to be used elsewhere.

Table 1: Module Input and Output File Types

| Module | Min. Input Files | Output Files |
|---|---|---|
| FASTAptameR-Count | Preprocessed FASTQ/A | FASTA or CSV |
| FASTAptameR-Translate | Counted FASTA | FASTA or CSV |
| FASTAptameR-Motif_Search | Counted FASTA | FASTA or CSV |
| FASTAptameR-Motif_Tracker | 2+ counted FASTAs | CSV |
| FASTAptameR-Distance | Counted FASTA | FASTA or CSV |
| FASTAptameR-Enrich | 2+ counted FASTAs | CSV |
| FASTAptameR-Positional_Enrichment | Enrich CSV | Plots |
| FASTAptameR-Cluster | Counted FASTA | FASTA or CSV |
| FASTAptameR-Cluster_Diversity | Clustered FASTA | CSV |
| FASTAptameR-Cluster_Enrich | 2+ CSVs from Cluster_Diversity | CSV |

#### 1.2 A note on plotting in FASTAptameR 2.0

Though many plots are initially created with `ggplot2`, they are all shown as interactive `plotly` plots. As such, you will see a number of options appear along the top of the image in response to your mouse hover.

These options will allow you to zoom in and out, select regions of interest, and download the plots. This last functionality is provided by the camera icon (first icon on the left as you hover over the image of the plots). Finally, double-clicking the image should reset it (*e.g.*, remove zoom or crop effects).

### 2 How to get started

Like its predecessor FASTAptamer, FASTAptameR 2.0 is designed to be easy to use and accessible for practitioners of combinatorial selection, and to be open-source so that the community can interact with the source code. There are three ways for users to interact with FASTAptameR 2.0. The *web server* (https://fastaptamer2.missouri.edu/) is the easiest way to interact with this application because it only requires an internet connection and browser. The user interface can also run on your local system as a *Docker container*. Finally, all code can be pulled from GitHub at https://github.com/SkylerKramer/FASTAptameR-2.0.

#### 2.1 Web server interface

The web server interface is the fastest and easiest way to use the FASTAptameR 2.0 User Interface, especially for relatively small data sets. The web server can be accessed from https://fastaptamer2.missouri.edu/, which is hosted by the Digital Biology Laboratory under the direction of Dr. Dong Xu at the University of Missouri - Columbia. However, this option only works if the files uploaded to any given module are less than 2 GB.

#### 2.2 Docker interface

The web server can also be run locally on your machine(s) via Docker. This interface functions identically to the web server interface and allows the user full access to all FASTAptameR 2.0 functions without having to rely on continuous internet connection and without the limitations of files sizes or file transfer speeds. In many cases, using the Docker interface is the most reliable and convenient way to explore your data set.

Docker is a convenient tool that may be used to construct *images* of software. The *image* functions as the blueprint for an application. The *image* of FASTAptameR 2.0, for example, contains all relevant software (*e.g.*, `R`), files (*e.g.*, this PDF), and packages (*e.g.*, `Shiny`).

The **three steps** required for initial installation of FASTAptameR 2.0 via Docker. **Step 1** is required for the initial installation of Docker. **Step 2** is required to get access to FASTAptameR 2.0 and all subsequent versions. **Step 3** is required to run the application. After initial installation of a given version of FASTAptameR 2.0, only Step 3 needs to be repeated in subsequent sessions. Instructions for accessing newer versions as they become available are in the instructions below for Step 2.

**Step 1. Install Docker to establish a "Docker-active terminal" on your local system.**

The first step is either to install Docker (on a Linux machine) or to install Docker Desktop (on a Windows or Mac machine) to establish a "Docker-active terminal" on your local system. For reference, Windows users can access their terminal (the Command Prompt) by searching the machine for `cmd`. Mac users can access their terminal by searching for `Terminal` from the Launchpad, Finder, Spotlight, *etc.*

If you would like to see extensive details on Docker or its installation, please visit https://www.docker.com/ and https://docs.docker.com/get-docker/, respectively. Importantly, the FASTAptameR 2.0 *image* is built on Linux. Thus, it is necessary to run it from a Linux environment or virtual machine. This should automatically happen after installing Docker for Linux or Docker Desktop for Mac. Windows users must also install Docker Desktop and will typically need to enable virtualization through their Bios (see https://www.tutorialspoint.com/windows10/windows10_virtualization.htm for more details).

Successful completion of this step will yield a "Docker-active terminal". Linux, Windows, or Mac users can check if their terminal is Docker-active by running the following command in the terminal, which should return which version of Docker has been installed if it was installed successfully:

```
docker version
```

**Step 2. Pull the FASTAptameR 2.0 image from the Docker Hub repository.**

The FASTAptameR 2.0 *image* must be pulled from a repository (*i.e.*, Docker Hub) by running the following command in a Docker-active terminal:

```
docker pull skylerkramer/fastaptamer2:publicupload05
```

Please note that `publicupload05` is the most recent version of FASTAptameR 2.0 as of the time of writing this tutorial. To access future updates as they become available, please refer to this section of the dynamic User Guide at https://github.com/SkylerKramer/FASTAptameR-2.0 or at the main Docker Hub page at https://hub.docker.com/repository/docker/skylerkramer/fastaptamer2.

**Step 3. Launch FASTAptameR 2.0.**

Once you have this application's image, run the following from a Docker-active terminal:

```
docker run -d --rm -p 3838:3838 skylerkramer/fastaptamer2:publicupload05
```

This will launch a local instance - a *container* - of FASTAptameR 2.0. You will then interact with this container in the same fashion as the web server by entering `localhost:3838` into your web browser address bar.

**Explanation of flags from Step 3:**

- `-d`: enable detached mode, which allows you to use your command line/terminal even with the active *container* (*i.e.*, *container* is detached from your terminal and runs in the background)
- `--rm`: automatically stop the container upon exit
- `-p 3838:3838`: publish `3838` host port (first number) to the `3838` container port (second number)
- `skylerkramer/fastaptamer2:publicupload05`: the local path to the **FASTAptameR 2.0** Docker *image*; please see **Step 2** for the most recent version
- `localhost:3838`: navigate here from your web browser to start interacting with **FASTAptameR 2.0**

#### 2.3 R interface

The third way to interact with FASTAptameR 2.0 is through the source code. All code is publicly available at https://github.com/SkylerKramer/FASTAptameR-2.0. This will allow R developers to adjust the code to their specific needs. Please note that all dependencies in this app (*e.g.*, Shiny, ggplot2, *etc.*) must be installed to use this app. A full list of dependencies and their installation instructions are available on the GitHub page above.

#### 2.4 Software usage

If you use, adapt, or modify FASTAptameR 2.0, please cite: (Alam KK 2015) and ("FASTAptameR 2.0," n.d.).

For any questions or concerns, please or make a GitHub issue.

### 3 Tutorial

#### 3.1 Data requirements

FASTAptameR 2.0 utilizes many string-based functions. Thus, this program can be used to analyze many types of biological populations. However, all libraries must be initially saved in a FASTA or FASTQ format and passed through **FASTAptameR-Count** prior to any subsequent analyses. Further, any data preprocessing steps, such as trimming flanking sequences or filtering for read quality, must be made outside of this application.

#### 3.2 Sample Data and Uploading User Data

##### 3.2.1 Sample data

All data shown in this tutorial come from published selections for RNA aptamers with affinity for HIV-1 reverse transcriptase (RT).

**Rnd14**: Briefly, 14 rounds of selection gave a population that was dominated by pseudoknots, especially those with a well-defined motif known as the Family 1 Pseudoknot (F1Pk) (Burke DH 1996).

**Rnd15**: Later analysis monitored population dynamics in response to increasing selection pressure and revealed two other RNA motifs - the '(6/5) asymmetric loop' and the 'UCAA' motif - that are more potent inhibitors of RT than the previously identified pseudoknots (Ditzler MA 2013; Whatley AS 2013). Notably, RT inhibition of F1Pk aptamers was abolished by the R277L mutation while '(6/5) asymmetric loop' and the 'UCAA' motif aptamers were insensitive to this mutation.

**Rnd17**: We then subjected the Rnd14 population above to three additional cycles of selection for affinity to RTs from phylogenetically diverse lentiviruses, including several that carried the R227K mutation. This "Poly-Target" selection approach identified aptamer subsets with broad target recognition and RT inhibition (Alam et al. 2018).

All of these data are available as preprocessed (trimmed and filtered) FASTA files from http://burkelab.missouri.edu/fastaptamer.html.

##### 3.2.2 Moving on to your data

To start analyzing the sample data or your own data, please do one of two things. Either **1)** upload a local copy of the file via the file browser in **FASTAptameR-Count** or **2)** supply a link to the data via the text box labeled as `Online source` in **FASTAptameR-Count**. This module is the entry point to **FASTAptameR 2.0**, so each analysis should start here.

#### 3.3 FASTAptameR-Count

##### 3.3.1 Description

Analyzing sequences from combinatorial selections typically begins with counting how many of each species are present. This baseline information immediately establishes whether the population has converged on a small handful of dominant sequences or on a large number of enriched sequences, of if instead there is still an enormous diversity with little convergence. Each of these scenarios is immediately evident in a plot of the total number of reads ordered by rank. Similarly, a simple count of how many of a given species is present is the input data required for many other layers of analysis.

Thus, the pipeline begins with FASTAptameR-Count, which is the gateway to the rest of FASTAptameR 2.0. Sequence files must first be processed through the FASTAptameR-Count module to generate a new file that removes redundant copies and tabulates the abundance of each sequence, its rank within the population, and a normalized value of the abundance (reads per million, or 'RPM') that normalizes the raw number of reads to the population size. This normalization enables comparisons of relative abundances across populations of different sizes. These statistical values are written into the sequence identifiers of the downloadable FASTA file.

FASTAptameR-Count serves as the entry point into this suite of modules, and, thus, it should be run prior to any of the following modules. Keep the *count* file in an easily accessible folder, as this file will serve as input for many other FASTAptameR 2.0 modules (**Fig. 1**).

FASTAptameR-Count accepts either a FASTQ/A file chosen with the file browser or a link to such a file (*e.g.*, the default GitHub link in the text box labeled as `Online source`) and returns a *counted* data table as output that can be downloaded as a FASTA or CSV file.

Input FASTQ files should be properly formatted (4 lines per entry with the second line of each entry being the sequence). Input FASTA files are not required to have sequence identifiers. No pre-existing sequence identifiers will be conserved by this module. Instead, output sequence identifiers are defined by the statistical representation of each sequence (specifically, the rank order of that sequence in the population, its read count, and its normalized read count - see below). Sample input files are shown in **Fig. 2**.

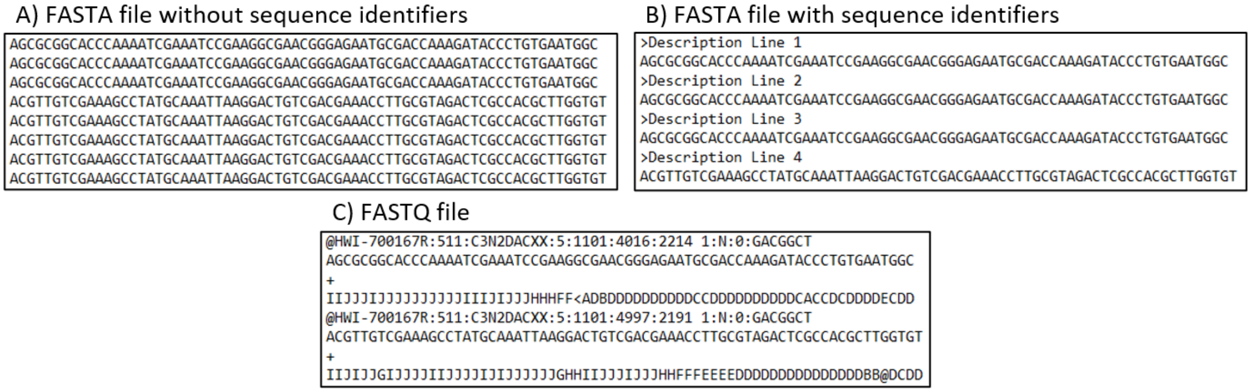

Figure 2: Examples of valid inputs to FASTAptameR-Count. These are all valid inputs to FASTAptameR-Count, which is the first step in using this workflow. A) FASTA file without sequence identifier lines. B) FASTA file with sequence identifier lines. C) FASTQ file.

All modules directly connected to FASTAptameR-Count are shown in **Fig. 3**, and a screenshot of the module interface is shown in **Fig. 4**.

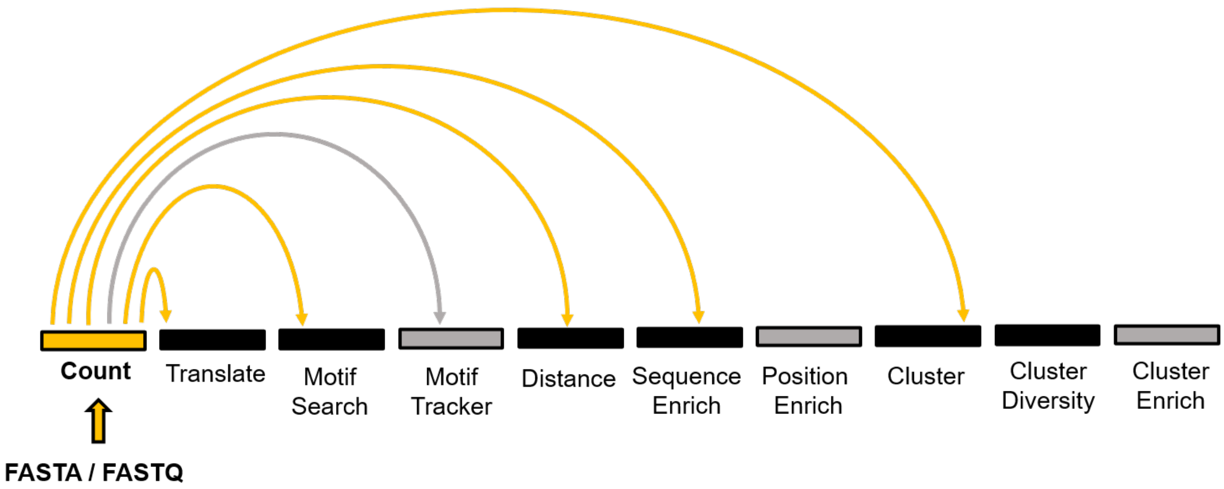

Figure 3: All modules connected to FASTAptameR-Count.

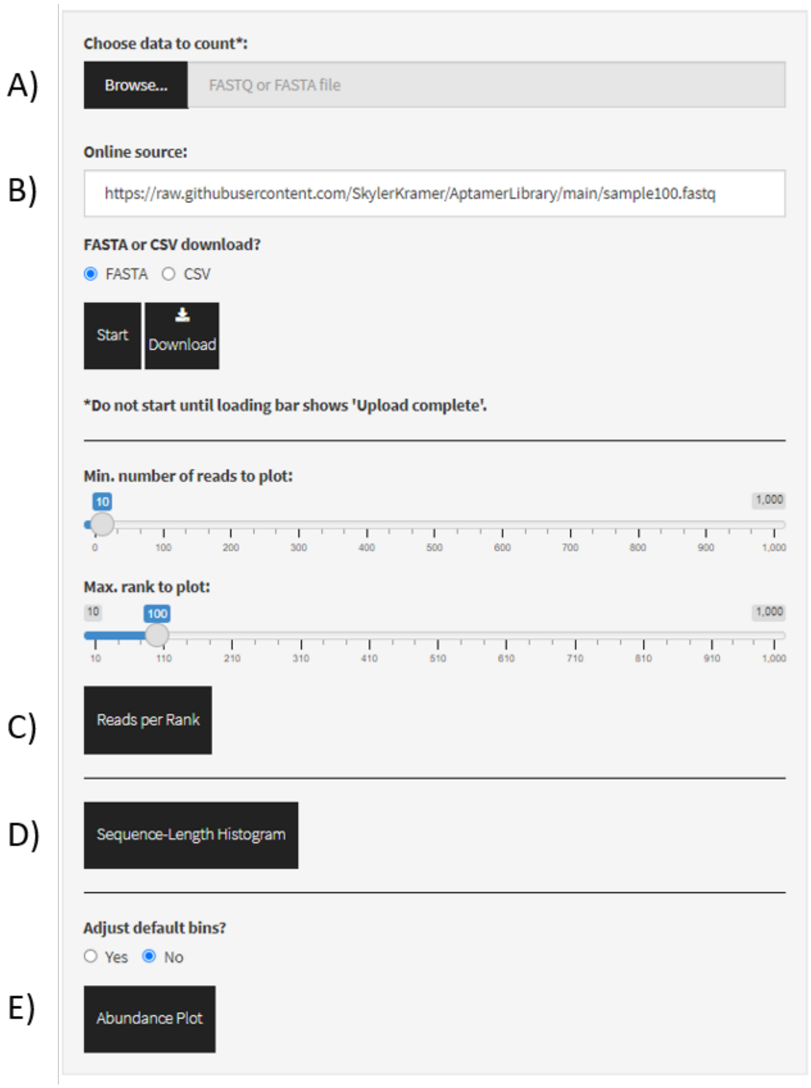

Figure 4: Screenshot of FASTAptameR-Count.

##### 3.3.2 Usage

The input FASTQ/A file must be chosen with the file browser (**Fig. 4A**) or linked in the text box (**Fig. 4B**). A sample link is already provided in the text box. Note, if a file is uploaded via the file browser **AND** a link is provided, only the uploaded file (**NOT** the linked one) will be analyzed.

For file uploads, please wait for the loading bar to show *Upload complete* before using the Start button. Once the files are fully uploaded or are chosen via the file server, the `Start` button will begin the counting process. The results will be displayed as a data table on the right side of the screen.

A sample output data table is shown in **Fig. 5**. Importantly, numeric columns in this output table and all other output tables are filterable by range (*e.g.*, to select the top 100 ranked sequences, type `1 ... 100` into the `Rank` column). To use some later features of FASTAptameR 2.0 (*i.e.*, the Distance and Position Enrichment modules), you must apply such a filter to the `Length` column such that all sequences are of the same length (*e.g.*, `70 ... 70` only retains sequences of length 70).

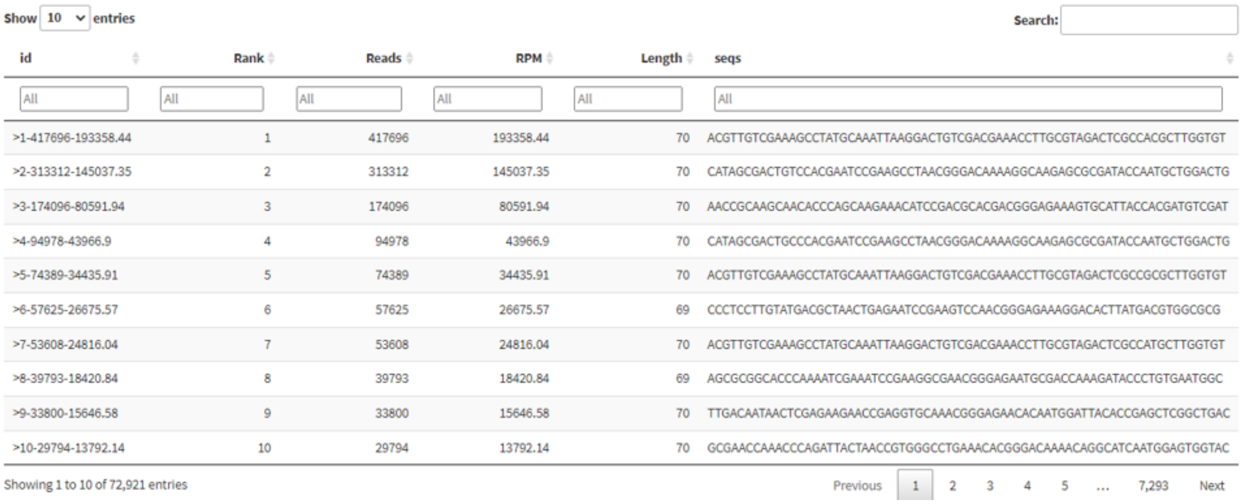
Show 10 entries  Search:

| id | Rank | Reads | RPM | Length | seqs |
|---|---|---|---|---|---|
| All | All | All | All | All | All |
| >1-417696-193358.44 | 1 | 417696 | 193358.44 | 70 | ACGTTGTCGAAAGCCTATGCAAATTAAGGACTGTCGACGAAACCTTGCGTAGACTCGCCACGCTTGGTGT |
| >2-313312-145037.35 | 2 | 313312 | 145037.35 | 70 | CATAGCGACTGTCCACGAATCCGAAGCCTAACGGGACAAAAGGCAAGAGCGCGATACCAATGCTGGACTG |
| >3-174096-80591.94 | 3 | 174096 | 80591.94 | 70 | AACCGCAAGCAACACCCAGCAAGAAACATCCGACGCACGACGGGAGAAAGTGCATTACCACGATGTCGAT |
| >4-94978-43966.9 | 4 | 94978 | 43966.9 | 70 | CATAGCGACTGCCCACGAATCCGAAGCCTAACGGGACAAAAGGCAAGAGCGCGATACCAATGCTGGACTG |
| >5-74389-34435.91 | 5 | 74389 | 34435.91 | 70 | ACGTTGTCGAAAGCCTATGCAAATTAAGGACTGTCGACGAAACCTTGCGTAGACTCGCCGCGCTTGGTGT |
| >6-57625-26675.57 | 6 | 57625 | 26675.57 | 69 | CCCTCCTTGTATGACGCTAACTGAGAATCCGAAGTCCAACGGGAGAAAGGACACTTATGACGTGGCGCG |
| >7-53608-24816.04 | 7 | 53608 | 24816.04 | 70 | ACGTTGTCGAAAGCCTATGCAAATTAAGGACTGTCGACGAAACCTTGCGTAGACTCGCCATGCTTGGTGT |
| >8-39793-18420.84 | 8 | 39793 | 18420.84 | 69 | AGCGCGGCACCCAAAATCGAAATCCGAAGGCGAACGGGAGAATGCGACCAAAGATACCCTGTGAATGGC |
| >9-33800-15646.58 | 9 | 33800 | 15646.58 | 70 | TTGACAATAACTCGAGAAGAACCGAGGTGCAAACGGGAGAACACAATGGATTACACCGAGCTCGGCTGAC |
| >10-29794-13792.14 | 10 | 29794 | 13792.14 | 70 | GCGAACCAAACCCAGATTACTAACCGTGGGCCTGAAACACGGGACAAAACAGGCATCAATGGAGTGGTAC |

Showing 1 to 10 of 72,921 entries  Previous 1 2 3 4 5 … 7,293 Next

Figure 5: FASTAptameR-Count Output. Note the search box in the top right corner. This appears in most module outputs and allows the user to search the table for specific strings.

Note that the *id* column has the following format: `>Rank-Reads-RPM`, where `Rank` is the order of sequences after sorting by `Reads`, which is the raw abundance of each sequence. `RPM` (*i.e.*, Reads per Million) is the value of `Reads`, normalized by the total population size: `RPM = Reads / (Population Size / 1e6)`.

The total number of sequences, number of unique sequences, and module runtime will be displayed below the `Start` and `Download` buttons after running is finished. The `Download` button opens a file browser prior to downloading the output as a FASTA or CSV file (`DEFAULT = FASTA`, which is required for subsequent modules). Again, keep the *count* file in an easily accessible folder, as this file will serve as input for many other FASTAptameR 2.0 modules.

##### 3.3.3 Troubleshooting

Do not start the module before the upload is finished. If you start the module before the upload is finished **AND** a file link is provided, then the *file link will be analyzed.* If you start the module before the upload is finished **AND** no link is provided, then you will *get an error message* that says `No file or link provided!`. If you start the module before the upload is finished **AND** no file link is provided **AND** you have previously uploaded a file to this module, then the *previously uploaded file will be reanalyzed.* In any of these cases, reuploading the file, *waiting for it to finish uploading*, and then starting the module should correctly analyze your data. If any errors persist, please refresh the page. Examples of incomplete and complete progress bar displays are given in **Fig. 6**.

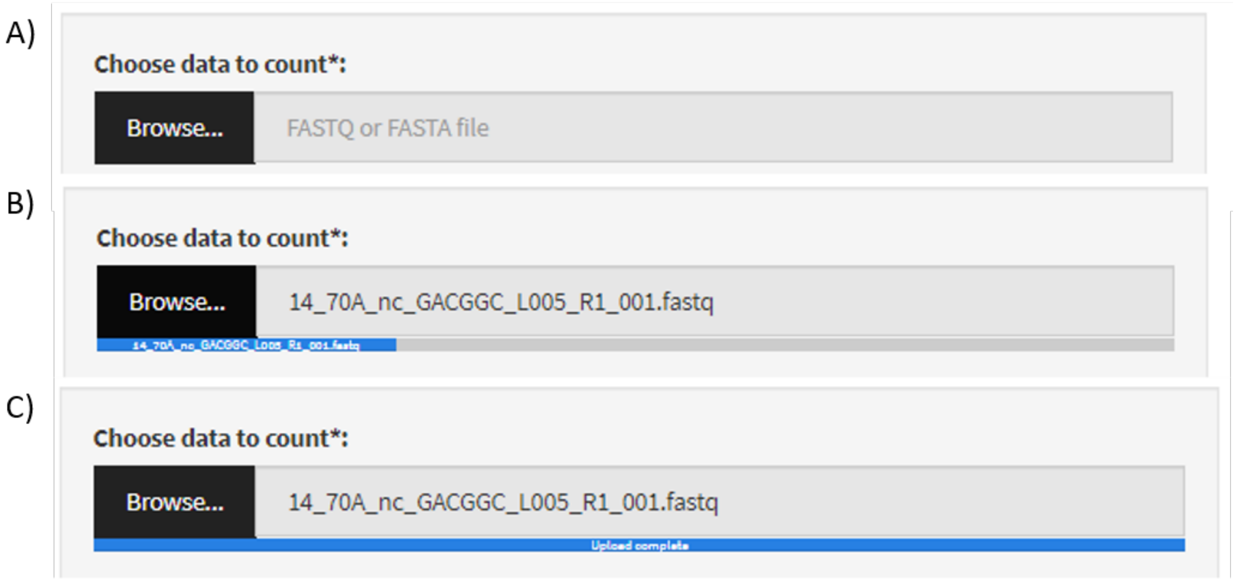

Figure 6: Example loading bars. A) Widget for file selection before upload. B) Loading bar (blue) during upload. C) Loading bar after successful upload. In any module requiring a file upload (*i.e.*, every module except for FASTAptameR-Count when a link is provided), this loading bar **must** show `Upload complete` before pressing the start button.

##### 3.3.4 Plotting

This module can generate three types of interactive plots based on the counted data to illustrate the overall distribution of sequence abundances in the population: a line plot of reads for each unique sequence sorted by rank (button shown in **Fig. 4C**, output shown in **Fig. 7A, B**), two histograms of sequence lengths (button shown in **Fig. 4D**, output shown in **Fig. 7C**), and a sequence abundance bar plot (button shown in **Fig. 4E**, output shown in **Fig. 7D**). Line plots are filterable by 1) minimum number of reads to plot and 2) maximum rank to plot, and both values are chosen with a slider bar. The histograms, however, are not filterable.

The sequence abundance bar plot first bins sequences based on their read counts and then plots these bins against their relative abundance (as fractions of the total population). Finally, the bars are colored according to the number of unique sequences in each bin. The breakpoints forming these bins are, by default, set to the following: `Reads = 1`, `1 < Reads < 10`, `10 <= Reads < 100`, `100 <= Reads < 1000`, `1000 <= Reads <= max(Reads)`. However, the user can toggle singletons on/off or customize these breakpoints by selecting `Yes` in the respective prompt. New breakpoints should be entered as a comma-separated list. An example is given in **Fig. 8**.

The first bin contains all sequences with `Reads < min(break point)` unless singletons are desired, in which case the set of sequences with `Reads = 1` becomes its own respective bin. The final bin contains all sequences with `max(break point) <= Reads`. Intermediate bins contain sequences between consecutive breakpoints, where the minimum and maximum values are inclusive and exclusive, respectively. For example, if singletons are not desired and the breakpoints are `20,2000`, then the bins are as follows: (`Reads < 20`), (`20 <= Reads < 2000`), and (`2000 <= Reads <= max(Reads)`).

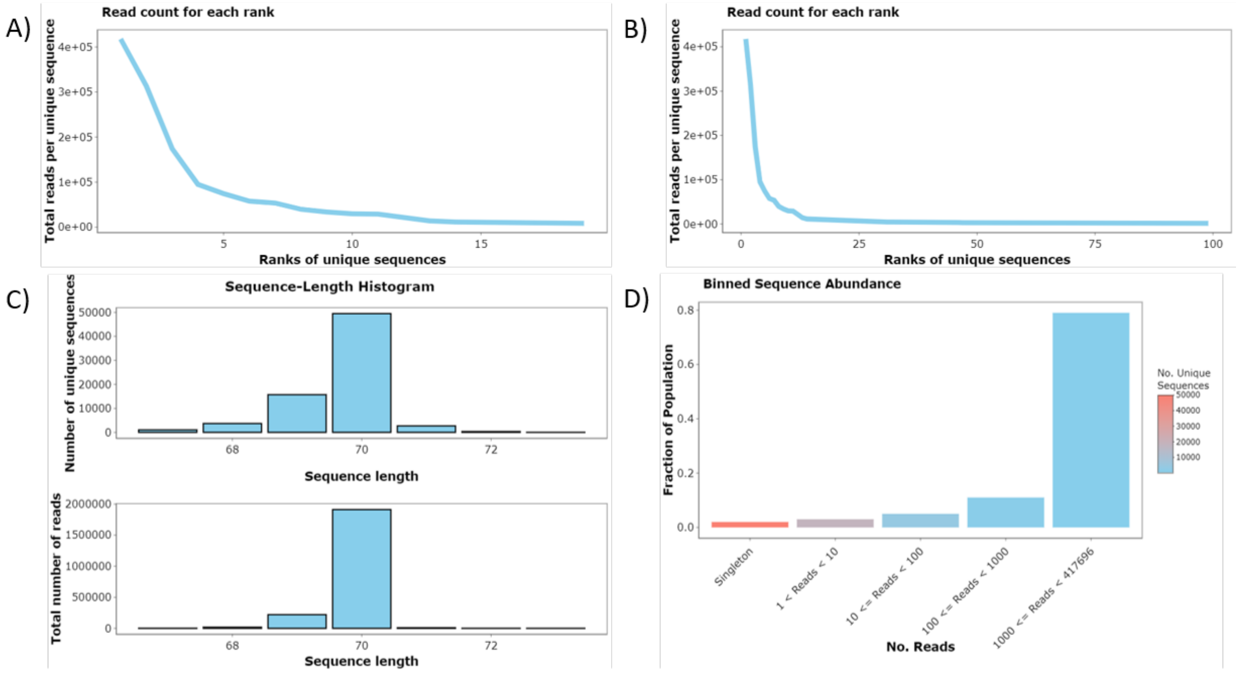

Figure 7: FASTAptameR-Count Plots. A) A line plot showing the total number of reads for the 20 most abundant sequences. B) The same plot with the 100 most abundant sequences. C) Histograms of sequence lengths for unique sequences (top) and all reads (bottom). D) Binned sequence abundance bar plot where `x` corresponds to discrete bins of read counts, `y` corresponds to the fraction of the total population, and color corresponds to the number of unique sequences per bin.

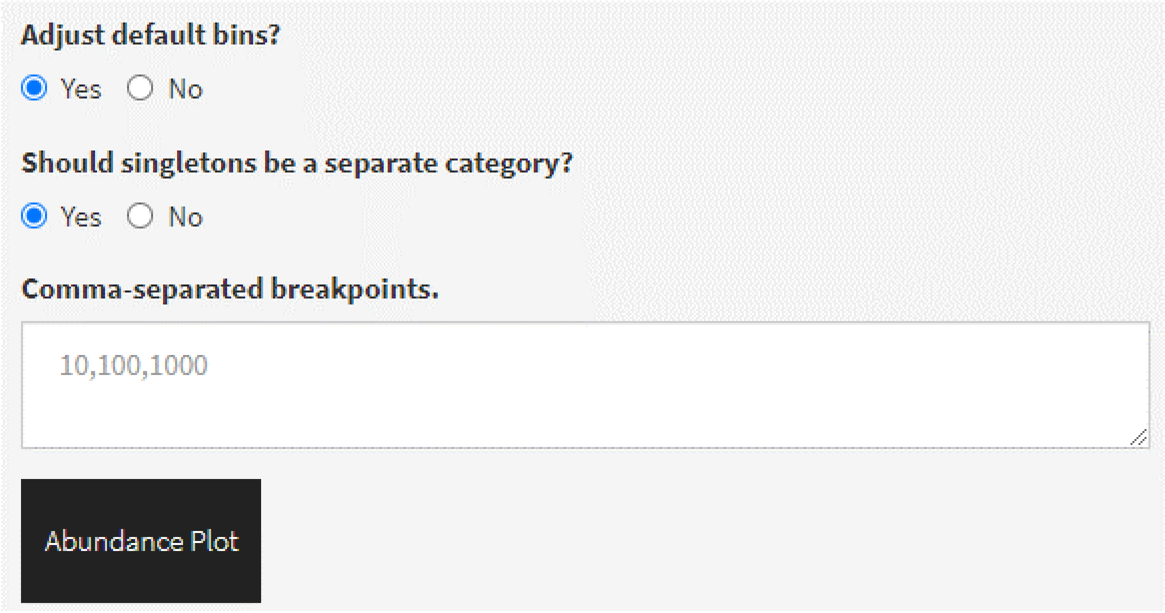

Figure 8: Example of how to customize the bins of the sequence abundance plot. Top radio button must be set to `Yes`. The second radio button determines whether singletons are a separate bin. The text input takes a comma-separated list of breakpoints. This image reflects the default breakpoints (listed in the text).

#### 3.4 FASTAptameR-Translate

##### 3.4.1 Description

In addition to analyzing nucleic acid populations, FASTAptameR 2.0 can also analyze combinatorial libraries based on translated peptide or protein products such as phage display, partially randomized cellular proteins selected for bioactivity, and evolving viral populations from patients. In addition, protein engineering and synthetic biology research makes increasing use of non-standard amino acids and expanded genetic alphabets; both of these innovations can be analyzed with FASTAptameR 2.0.

FASTAptameR-Translate translates input nucleotide sequences into amino acid sequences following either the standard genetic code, a biologically derived, nonstandard code, or a user-defined code. The input nucleotide sequences are treated as positive-sense mRNA. This module accepts a *counted* FASTA file and returns a *translated* data table that can be downloaded as a FASTA or CSV file.

All modules directly connected to FASTAptameR-Translate are shown in **Fig. 9**, and a screenshot of the module interface is shown in **Fig. 10**.

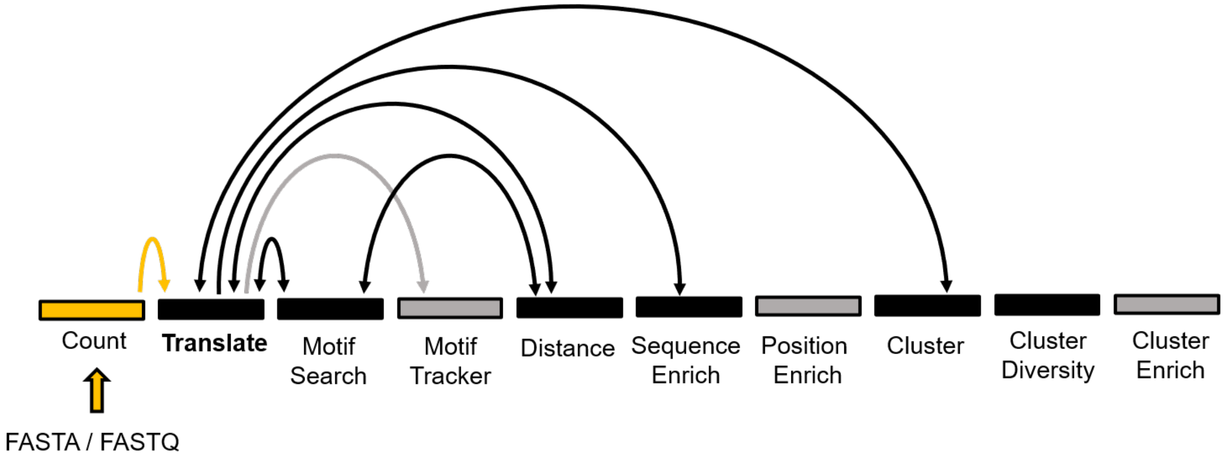

Figure 9: All modules connected to FASTAptameR-Translate.

##### 3.4.2 Usage

The input FASTA file must be chosen with the file browser. The open reading frame may be selected by the first set of radio buttons (`DEFAULT = 1`) (**Fig. 10A**). The second set of radio buttons indicates whether data for nucleotide sequences that encode the same amino acid sequence should be merged (`DEFAULT = Yes`) (**Fig. 10B**). If `Yes`, then redundantly encoded amino acid sequences are converged, and a new column (`Unique.Nt.Count`) will specify how many non-unique nucleotide sequences from the *counted* input were merged into each amino acid sequence. If `No`, then each unique nucleotide sequence is treated separately, even if multiple sequences encode the same amino acid sequence.

Merging the data for redundant sequences (button set to `No`) focuses attention on the encoded peptide sequences, irrespective of which nucleotide sequences gave rise to them, which can be especially informative for protein structure/function studies. In contrast, treating each evolving species separately (button set to `Yes`) allows users to discern contributions of nucleotide sequence to overall fitness by comparing relative enrichments of different sequences that that encode the same translated product.

The dropdown menu in this UI (**Fig. 10C**) allows the user to select one of sixteen genetic codes for translation (`DEFAULT = Standard`) (Gasteiger 2003):

1. Standard

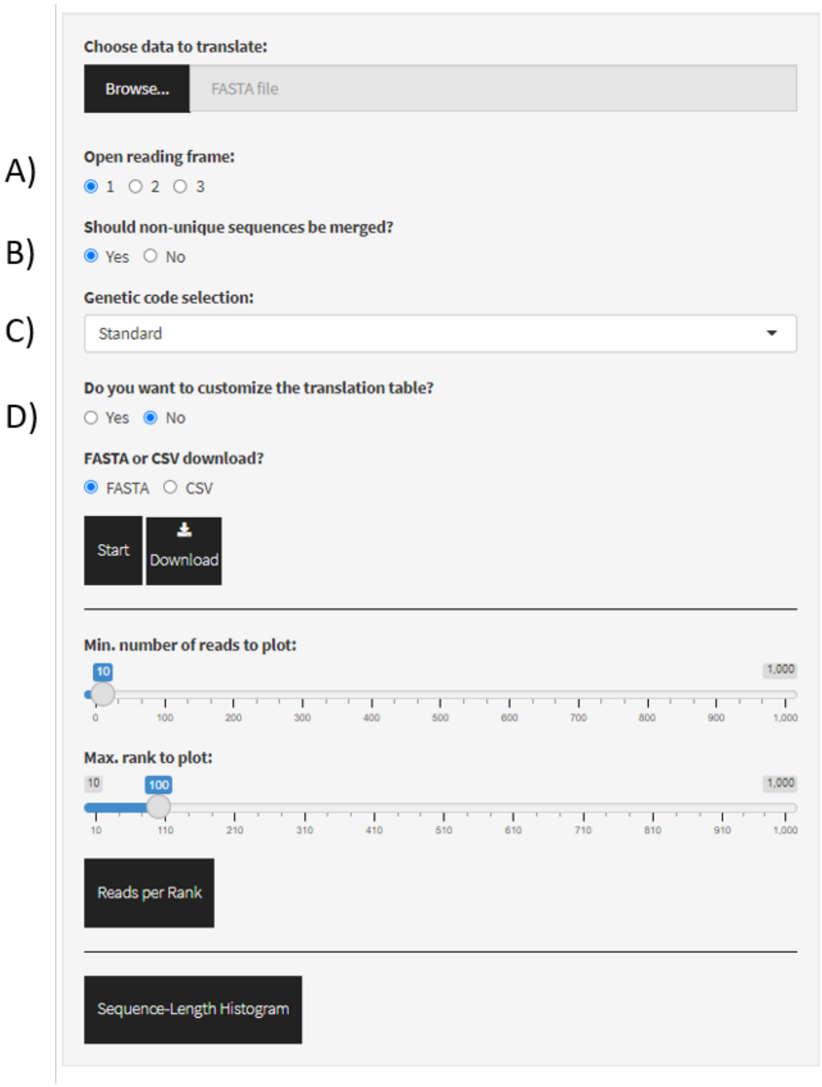

Figure 10: Screenshot of FASTAptameR-Translate.

2. Vertebrate mitochondrial
3. Yeast mitochondrial
4. Mold, protozoan, and coelenterate mitochondrial + Mycoplasma / Spiroplasma
5. Invertebrate mitochondrial
6. Ciliate, dasycladacean and Hexamita nuclear
7. Echinoderm and flatworm mitochondrial
8. Euplotid nuclear
9. Alternative yeast nuclear
10. Ascidian mitochondrial
11. Alternative flatworm mitochondrial
12. Blepharisma nuclear
13. Chlorophycean mitochondrial
14. Trematode mitochondrial
15. Scenedesmus obliquus mitochondrial
16. Pterobranchia mitochondrial

The user may also customize the translation code by selecting `Yes` in the third set of radio buttons (**Fig. 10D**) prior to translating. If `Yes`, then comma-separated codon / translation pairs may be entered in the resulting text box (*e.g.*, `GAT,Q` to recode `GAT` from its standard `Glu (E)` to `Gln (Q)`. Additional pairs can be included but must be on separate lines. If the codon already exists in the standard genetic code, then the user-supplied mapping will take precedence. If the codon does not exist in the standard genetic code, then it will be added to it.

Only 3-letter codons and 1-letter translations are currently accepted.

Non-standard alphabets are allowed for both input and output. For example, to translate an input amber STOP codon (`UGA`) as a nonstandard amino acid designated as `2`, the resulting text box would be entered as `TGA,2`. Similarly, to translate a triplet that contains the AEGIS nucleotide nitropyridine (`Z`) in the first position of a glycine codon, with `Trp` as the resulting output, the resulting text box would be as `ZGG,W`.

The `Start` button begins the translation process. The *translated* data table will be shown on the right side of the screen. The `Download` button opens a file browser prior to downloading the output as a FASTA or CSV file (`DEFAULT = FASTA`, which is required for subsequent modules).

##### 3.4.3 Plotting

This module generates a line plot of reads for each unique sequence sorted by rank and two histograms of sequence lengths. These plots illustrate overall population structure for the translated products and are analogous to the corresponding plots from FASTAptameR-Count. See that section for more details.

#### 3.5 FASTAptameR-Motif_Search

##### 3.5.1 Description

It is often useful to find all occurrences of a given sequence motif within a population. Even when parts of the functional structure are not defined simply by sequence (such as generic base-paired helices), many functional biomolecules contain 'signature sequences' that make useful first-pass filters. The motif of interest may be fully contiguous, such as the 8 nucleotide apical loop of the bacteriophage T4 gene 43 translational operator (`AAUACUC`) (Craig Tuerk and Gold 1990), or it may be discontinuous, such as the 11 nucleotides of the (6/5) asymmetric loop aptamer (abbreviated as "(6/5)AL"), in which a stem is interrupted by an asymmetric internal loop with the sequence `ARCGUY` on one strand and `RARAC` on the other. For the (6/5)AL motif, both elements must be present within the same sequence for that sequence to form a functional (6/5)AL aptamer. (Note that the `A` in `ARCGUY` is typically the last base of the 5' constant region and is removed when trimming sequences).

FASTAptameR-Motif_Search identifies sequences that contain one or more user-specified sequence motifs, or 'patterns.' The module accepts a *counted* FASTA file and returns a *searched* data table that can be downloaded as a FASTA or CSV file. Sequences in the output must have at least one occurrence of each pattern or at least one occurrence of at least one pattern (see details below for the `partial match` radio button).

All modules directly connected to FASTAptameR-Motif_Search are shown in **Fig. 11**, and a screenshot of the module interface is shown in **Fig. 12**.

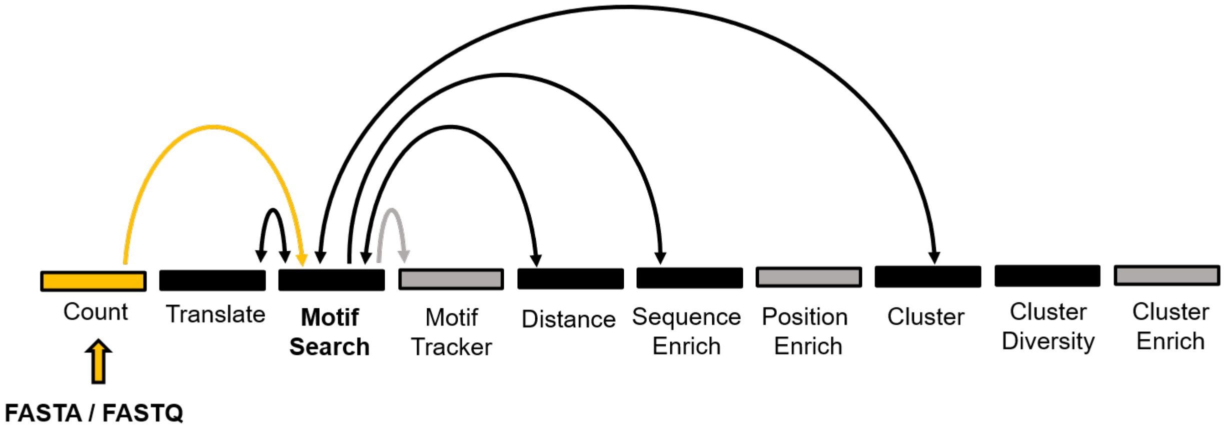

Figure 11: All modules connected to FASTAptameR-Motif_Search.

##### 3.5.2 Usage

The input FASTA file must be chosen with the file browser. The following text box (**Fig. 12A**) must contain at least one pattern (*e.g.*, `AAA`). If the user wishes to search for multiple patterns, the patterns must be separated by commas (*e.g.*, `AAA,GTG`).

The first set of radio buttons (**Fig. 12B**) determines whether the output has parentheses set around identified patterns. The default is to set to `No`, and patterns are highlighted in yellow in the output data without parentheses. For example, when `pattern = GGC` and `sequence = AAAGGCT`, the default output is `AAAGGCT` with `GGC` highlighted. Setting this button to `Yes` returns the output as `AAA(GGC)T`, and `GGC` is still highlighted. When two or more patterns overlap, output highlights the complete match when the button is set to `No` but only displays parentheses around the first queried search term that is matched when the button is set to `Yes`. For example, when `pattern = AGGC,GGCT` and `sequence = AAAGGCT`, the default output is `AAAGGCT` (with `AGGCT` highlighted), while it is `AA(AGGC)T` (with `AGGC` highlighted) when the parentheses

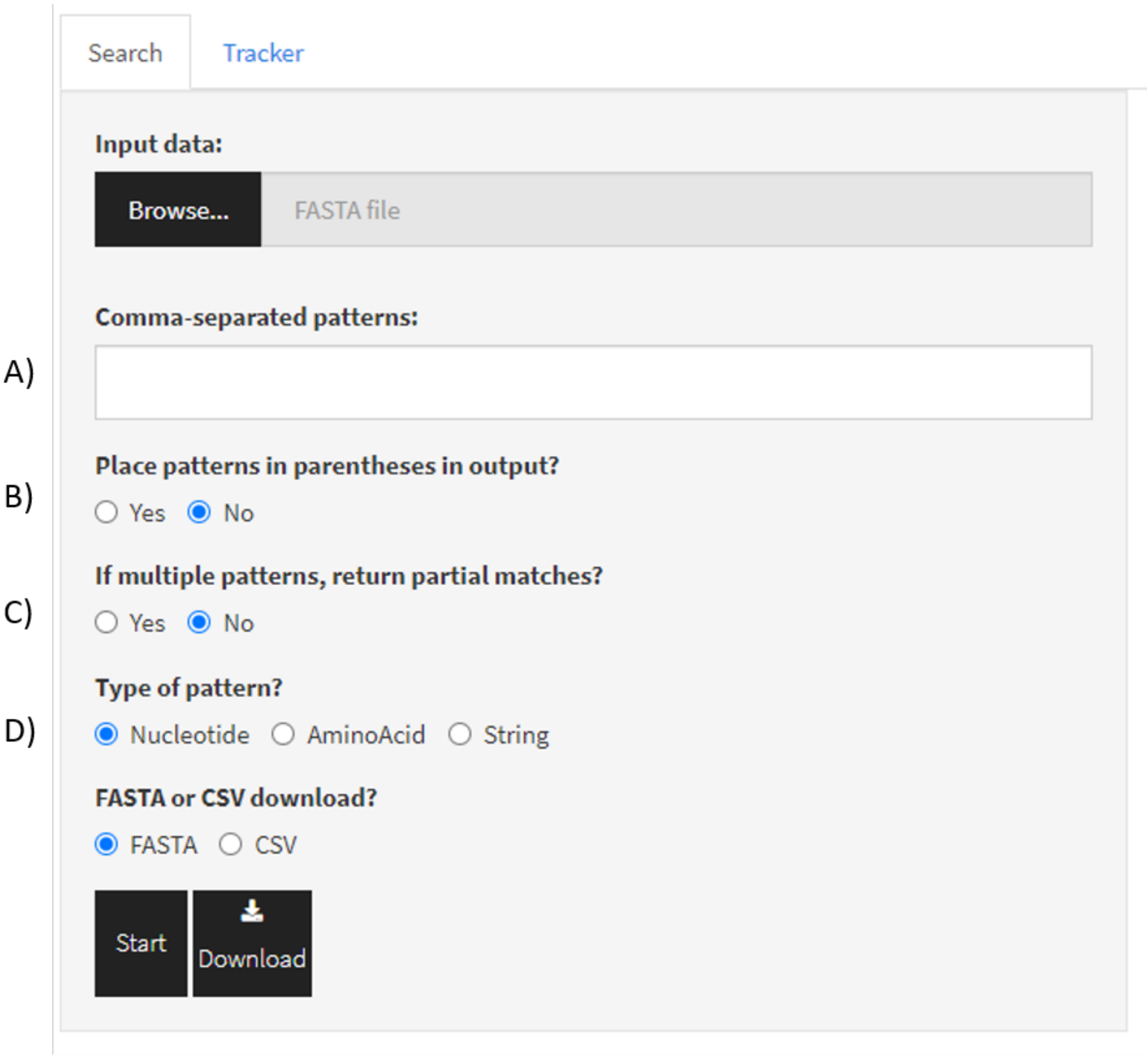

Figure 12: Screenshot of FASTAptameR-Motif_Search.

option is turned on. Note that parentheses will be treated as individual characters by subsequent modules and may alter downstream analyses.

The second set of radio buttons (**Fig. 12C**) governs how the software deals with multiple search terms. When the query contains multiple patterns, the search can be carried out either as a Boolean `AND` function by requiring all parts of the query to be present within a given sequence (this is the `default`, with button set to `No`), or as a Boolean `OR` function to identify sequences that contain any part of the query (set button to `Yes`). If `Yes`, filtered sequences must have at least one occurrence of **at least one** of the listed patterns. If `No` (`DEFAULT`), filtered sequences must have at least one occurrence of **each** of the listed patterns. For example, since both elements of the (6/5)AL aptamer must be present within the same sequence, the search for candidate (6/5)AL aptamers would be entered as `RCGUY,RARAC`, and this button would be set to `No`.

The third set of radio buttons (**Fig. 12D**) determines the type of pattern (`DEFAULT = Nucleotide`). If `Nucleotide`, then degenerate nucleotide codes are allowed, and T/U are interchangeable. Degenerate search patterns are **not** allowed for protein sequences or other sequence types. All patterns are converted to uppercase and have white spaces removed regardless of the pattern type.

1. **A/T/G/C/U** - single bases
2. **R** - puRine (A/G)
3. **Y** - pYrimidine (C/T)
4. **W** - Weak (A/T)
5. **S** - Strong (G/C)
6. **M** - aMino (A/C)
7. **K** - Keto (G/T)
8. **B** - not A
9. **D** - not C
10. **H** - not G
11. **V** - not T/U
12. **N** - aNy base (no *gap*)

The `Start` button begins the search process. The *searched* data table will be shown on the right side of the screen. The `Download` button opens a file browser prior to downloading the output as a FASTA or CSV file (`DEFAULT = FASTA`, which is required for subsequent modules).

A sample output data table is shown in **Fig. 13** with the following parameters: `comma-separated patterns = UCCG,CGGGAnAA`; `parentheses = No`; `partial filtering = No`; and `pattern type = Nucleotide`.

Show 10 entries    Search: TCCG|CGGGA[ACGT]AA

| id | Rank | Reads | RPM | seqs |
|---|---|---|---|---|
| All | All | All | All | All |
| >2-313312-145037.35 | 2 | 313312 | 145037.35 | CATAGCGACTGTCCACGAATCCGAAGCCTAACGGGACAAAAGGCAAGAGCGCGATACCAATGCTGGACTG |
| >3-174096-80591.94 | 3 | 174096 | 80591.94 | AACCGCAAGCAACACCCAGCAAGAAACATCCGACGCACGACGGGAGAAAGTGCATTACCACGATGTCGAT |
| >4-94978-43966.9 | 4 | 94978 | 43966.9 | CATAGCGACTGCCCACGAATCCGAAGCCTAACGGGACAAAAGGCAAGAGCGCGATACCAATGCTGGACTG |
| >6-57625-26675.57 | 6 | 57625 | 26675.57 | CCCTCCTTGTATGACGCTAACTGAGAATCCGAAGTCCAACGGGAGAAAGGACACTTATGACGTGGCGCG |
| >8-39793-18420.84 | 8 | 39793 | 18420.84 | AGCGCGGCACCCAAAATCGAAATCCGAAGGCGAACGGGAGAATGCGACCAAAGATACCCTGTGAATGGC |
| >12-22089-10225.37 | 12 | 22089 | 10225.37 | CATAGCGACTGTCCACGAATCCGAAGCCTAACGGGACAAAAGGCAGGAGCGCGATACCAATGCTGGACTG |
| >13-14115-6534.07 | 13 | 14115 | 6534.07 | CATAGCGACTATCCACGAATCCGAAGCCTAACGGGACAAAAGGCAAGAGCGCGATACCAATGCTGGACTG |
| >14-11313-5236.98 | 14 | 11313 | 5236.98 | AGCGCGGCACCCAAAATCGAAATCCGAAGGCGAACGGGAGAATGCGTCCAAAGATACCCTGTGAATGGC |
| >15-10818-5007.83 | 15 | 10818 | 5007.83 | CATAGCGACTGTCCACGAATCCGAAGCCTAACGGGACAAAAGGCAAGAGTGCGATACCAATGCTGGACTG |
| >16-10514-4867.11 | 16 | 10514 | 4867.11 | CATAGCGACCGTCCACGAATCCGAAGCCTAACGGGACAAAAGGCAAGAGCGCGATACCAATGCTGGACTG |

Showing 1 to 10 of 36,025 entries    Previous 1 2 3 4 5 … 3,603 Next

Figure 13: FASTAptameR-Motif_Search Output. Motifs are highlighted by default, and the specific patterns that get highlighted are shown in the search box. Omiting the search text from the search bar will remove the highlighting.

#### 3.6 FASTAptameR-Motif_Tracker

##### 3.6.1 Description

Individual species often rise and fall during combinatorial selections as a function of their overall fitness or other evolutionary forces, and they can enrich to different degrees in independently evolving populations. The same is true for sets of species that carry a given sequence motif. Tracking these changes across multiple populations can identify when certain species or sequence motifs first emerge (or disappear) and provides significant insights into their evolutionary dynamics.

FASTAptameR-Motif_Tracker reports on the occurrence of one or more query patterns / sequences across multiple populations. The module accepts at least two *counted* FASTA files as input and returns a data table of metadata related to the enrichment of the query pattern(s) across multiple populations. Multiple FASTA files should be selected from the file browser at the same time. Columns of the output data table include the following:

1. Population
2. File name
3. Sequences that contain the query
4. Rank
5. Reads
6. RPM

Optionally, an alias list of alternative motif/sequences names can be provided and will be included as a separate column. These aliases will be used in the legend of the line plot. If provided, there must be one alias per query per line (*e.g.*, `Seq1`, `Seq2`, `Seq3`; with each alias on its own line).

This module generates two data tables that can be downloaded as CSV files. The first table provides summary statistics for each query, and the second table provides enrichment scores for each query.

All modules directly connected to FASTAptameR-Motif_Tracker are shown in **Fig. 14**, and a screenshot of the module interface is shown in **Fig. 15**.

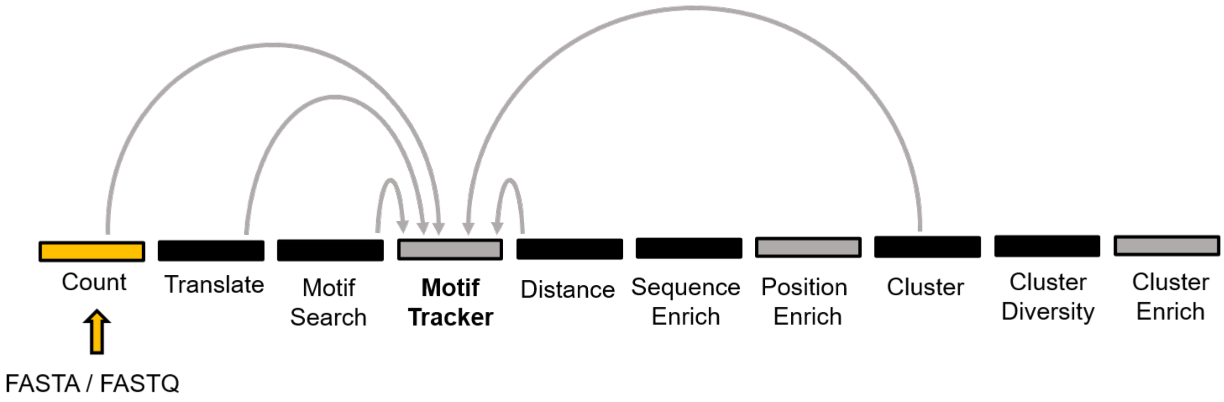

Figure 14: All modules connected to FASTAptameR-Motif_Tracker.

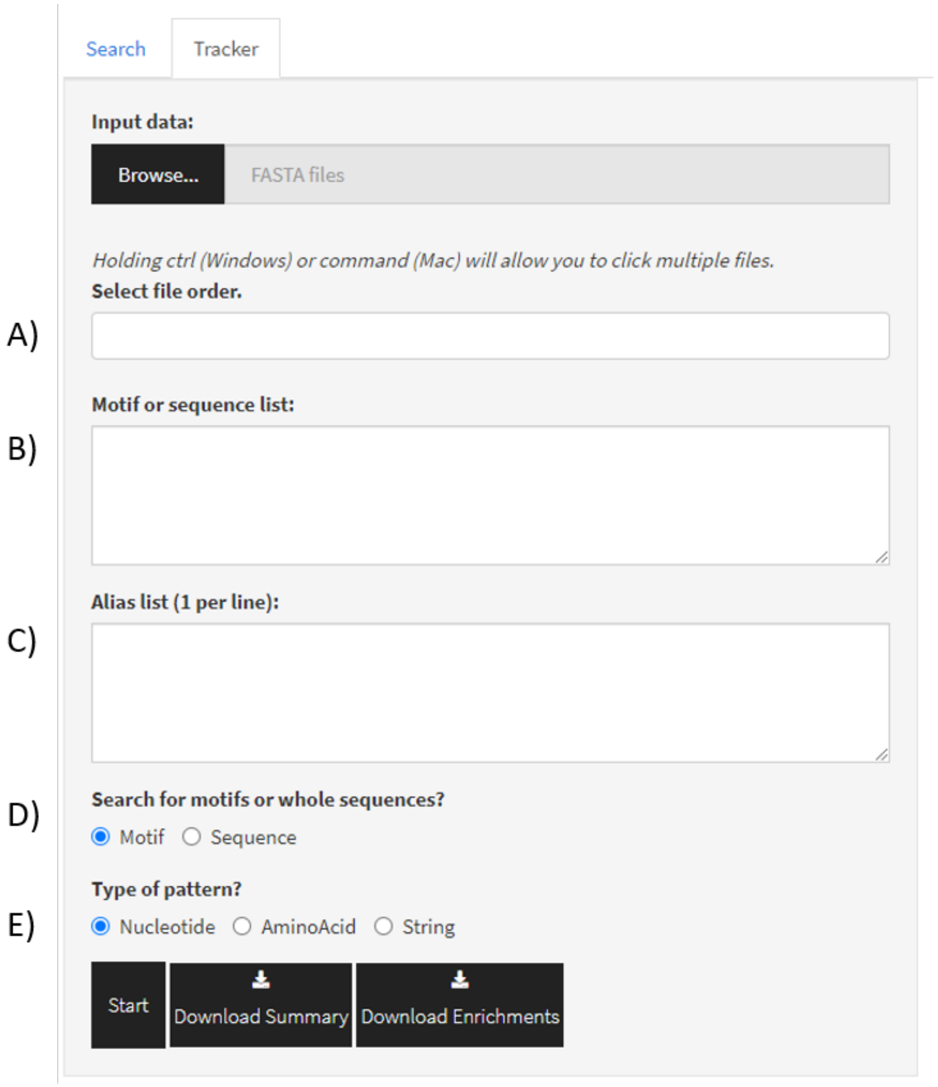

Figure 15: Screenshot of FASTAptameR-Motif_Tracker.

##### 3.6.2 Usage

The input FASTA files must be chosen with the file browser. The next line (**Fig. 15A**) determines the order of the input files via a dynamically generated dropdown menu. the order in which the user selects the files in this list determines which populations are considered to be population 1, which population 2, *etc.* The following text box (**Fig. 15B**) must contain at least one pattern or sequence. If the user wishes to search for multiple patterns or sequences, each pattern or sequence must be entered on separate lines. The text box for aliases **Fig. 15C** allows the user to rename the motifs or sequences to a more convenient name (such as 'Motif 1', 'G-rich', or 'F1Pk'), which will be used in the figure legend for ease of identification. Only one alias must be provided per line.

The first set of radio buttons (**Fig. 15D**) determines whether the queries are motifs or sequences, which determines how matches are identified. If `Motif`, then commas in the line are interpreted as separating submotifs, and regex pattern matching is used. For now, commas are interpreted as Boolean `AND` functions. If `Sequence`, then exact matches for each query are returned. The second set of radio buttons (**Fig. 15E**) determines the type of query (`DEFAULT = Nucleotide`). If `Nucleotide`, then degenerate nucleotide codes are allowed. Note, the query is converted to uppercase and white spaces are removed regardless of the type.

The `Start` button begins the motif enrichment process. The resulting data table will be shown on the right side of the screen. The `Download` button opens a file browser prior to downloading the output as a CSV file.

A screenshot of a sample output data table from tracked sequences is shown in **Fig. 16** with the three most abundant sequences from the 70HRT14 population as input queries. For the example search below, 70HRT14 is population 1 and 70HRT15 is population 2.

Show 10 entries Search:

| Population | FileName | seqs | Alias | Rank | Reads | RPM |
|---|---|---|---|---|---|---|
| 1 | 70HRT14-count.fasta | AACCGCAAGCAACACCCAGCAAGAAACATCCGACGCACGACGGGAGAAAGTGCATTACCACGATGTCGAT | 3rd | 3 | 174096 | 80591.94 |
| 2 | 70HRT15-count.fasta | AACCGCAAGCAACACCCAGCAAGAAACATCCGACGCACGACGGGAGAAAGTGCATTACCACGATGTCGAT | 3rd | 5 | 104932 | 52786.23 |
| 1 | 70HRT14-count.fasta | ACGTTGTCGAAAGCCTATGCAAATTAAGGACTGTCGACGAAACCTTGCGTAGACTCGCCACGCTTGGTGT | 1st | 1 | 417696 | 193358.44 |
| 2 | 70HRT15-count.fasta | ACGTTGTCGAAAGCCTATGCAAATTAAGGACTGTCGACGAAACCTTGCGTAGACTCGCCACGCTTGGTGT | 1st | 3 | 161830 | 81408.87 |
| 1 | 70HRT14-count.fasta | CATAGCGACTGTCCACGAATCCGAAGCCTAACGGGACAAAAGGCAAGAGCGCGATACCAATGCTGGACTG | 2nd | 2 | 313312 | 145037.35 |
| 2 | 70HRT15-count.fasta | CATAGCGACTGTCCACGAATCCGAAGCCTAACGGGACAAAAGGCAAGAGCGCGATACCAATGCTGGACTG | 2nd | 1 | 382391 | 192362.47 |

Showing 1 to 6 of 6 entries Previous 1 Next

Show 10 entries Search:

| Comparison | Query | Alias | Enrichment |
|---|---|---|---|
| 70HRT15-count.fasta : 70HRT14-count.fasta | AACCGCAAGCAACACCCAGCAAGAAACATCCGACGCACGACGGGAGAAAGTGCATTACCACGATGTCGAT | 3rd | 0.655 |
| 70HRT15-count.fasta : 70HRT14-count.fasta | ACGTTGTCGAAAGCCTATGCAAATTAAGGACTGTCGACGAAACCTTGCGTAGACTCGCCACGCTTGGTGT | 1st | 0.421 |
| 70HRT15-count.fasta : 70HRT14-count.fasta | CATAGCGACTGTCCACGAATCCGAAGCCTAACGGGACAAAAGGCAAGAGCGCGATACCAATGCTGGACTG | 2nd | 1.326 |

Showing 1 to 3 of 3 entries Previous 1 Next

Figure 16: FASTAptameR-Motif_Tracker Output.

##### 3.6.3 Plotting

This module can generate an interactive line plot showing the query's RPM across each population (**Fig. 17**). Although only two populations and three query sequences were used here for illustration, this tool can be especially useful for observing the rise and fall of multiple query sequences over the course of multiple rounds of selection.

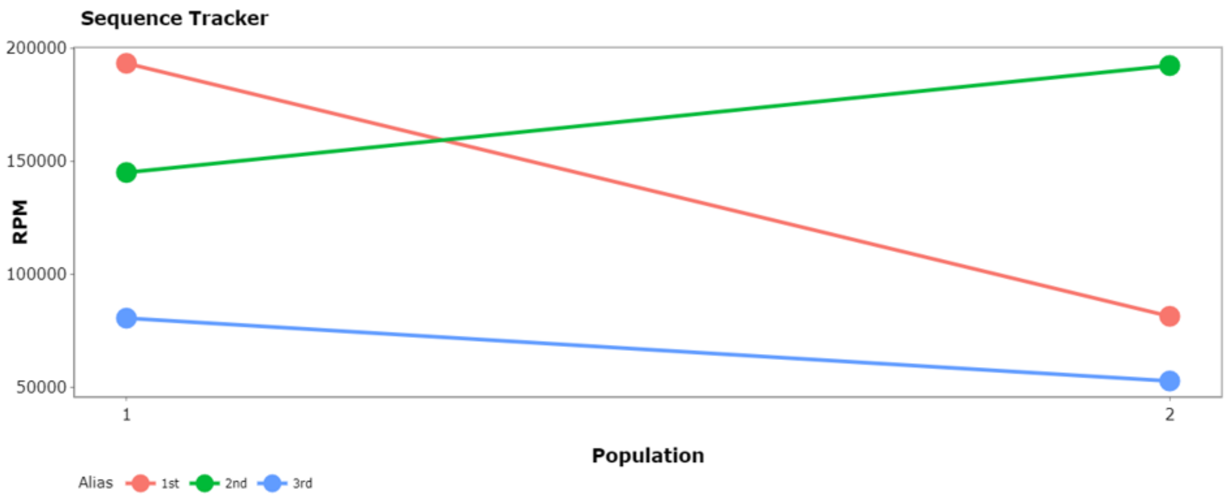

Figure 17: Sequence Tracking Line Plot. Shows the RPM of three sequences across the 70HRT14 and 70HRT15 populations. The aliases - ‘1st’, ‘2nd’, ‘3rd’ - refer to the first, second, and third most abundant sequences from the 70HRT14 population.

#### 3.7 FASTAptameR-Distance

##### 3.7.1 Description

The distribution of edit distances from a given “seed” sequence to the rest of the population informs population structure, evolutionary dynamics, and fitness landscapes. For example, a tight grouping of closely related sequences may reflect low mutation rates, rugged fitness landscapes, or strong purifying selective pressures. Alternatively, high diversity within a cluster may be the result of high mutation rates, smooth fitness landscapes, or weak selective pressures. A wide separation between that cluster and the rest of the population may indicate wide-spaced fitness peaks or sparsely sampled sequence space at the outset of the selection. In contrast, intermediate or closely spaced peaks may indicate that the starting population sampled sequence space more densely and that species selected for one fitness peak may be able to acquire the necessary mutations to sample other fitness peaks. Finally similar analyses can help to establish diversity or mutational density for a starting library prior to a selection.

FASTAptameR-Distance tabulates the distribution of distances from a user-defined reference sequence for all sequences in a population. The module accepts a *counted* FASTA file as input and returns a data table that contains a column for the Levenshtein edit distance (LED) between each input sequence and a query sequence. The LED is the minimum number of substitutions, insertions, or deletions required to transform one sequence into another. In that sense, it is more general than the Hamming distance, which only considers the minimum number of substitutions required to transform one sequence to another sequence of equal length. The output can be downloaded as a FASTA or CSV file.

All modules directly connected to FASTAptameR-Distance are shown in **Fig. 18**, and a screenshot of the module interface is shown in **Fig. 19**.

##### 3.7.2 Usage

The input FASTA file must be chosen with the file browser, and the following text box must contain a single query sequence. Note, this query sequence may not have any degenerate nucleotide codes. The `Start` button begins the distance calculations. The resulting data table will be shown on the right side of the screen. The `Download` button opens a file browser prior to downloading the output as a FASTA or CSV file.

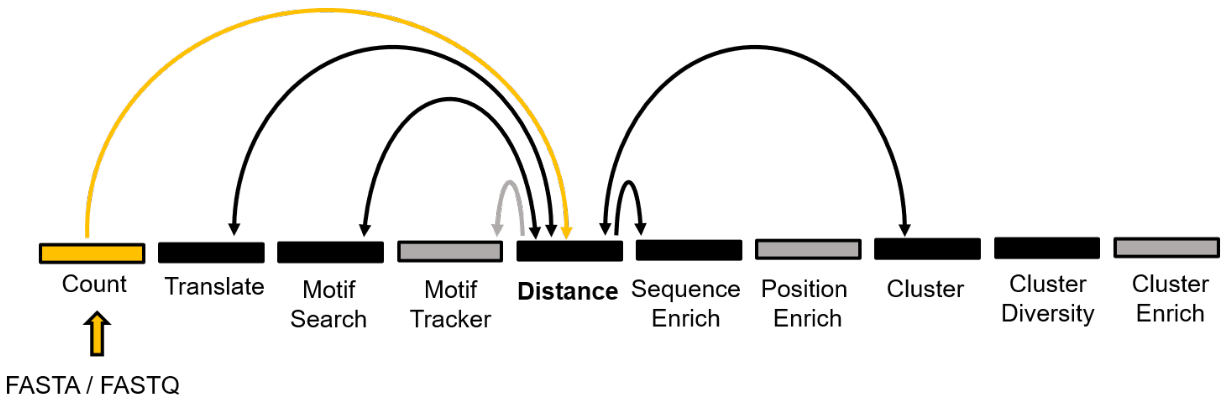

Figure 18: All modules connected to FASTAptameR-Distance.

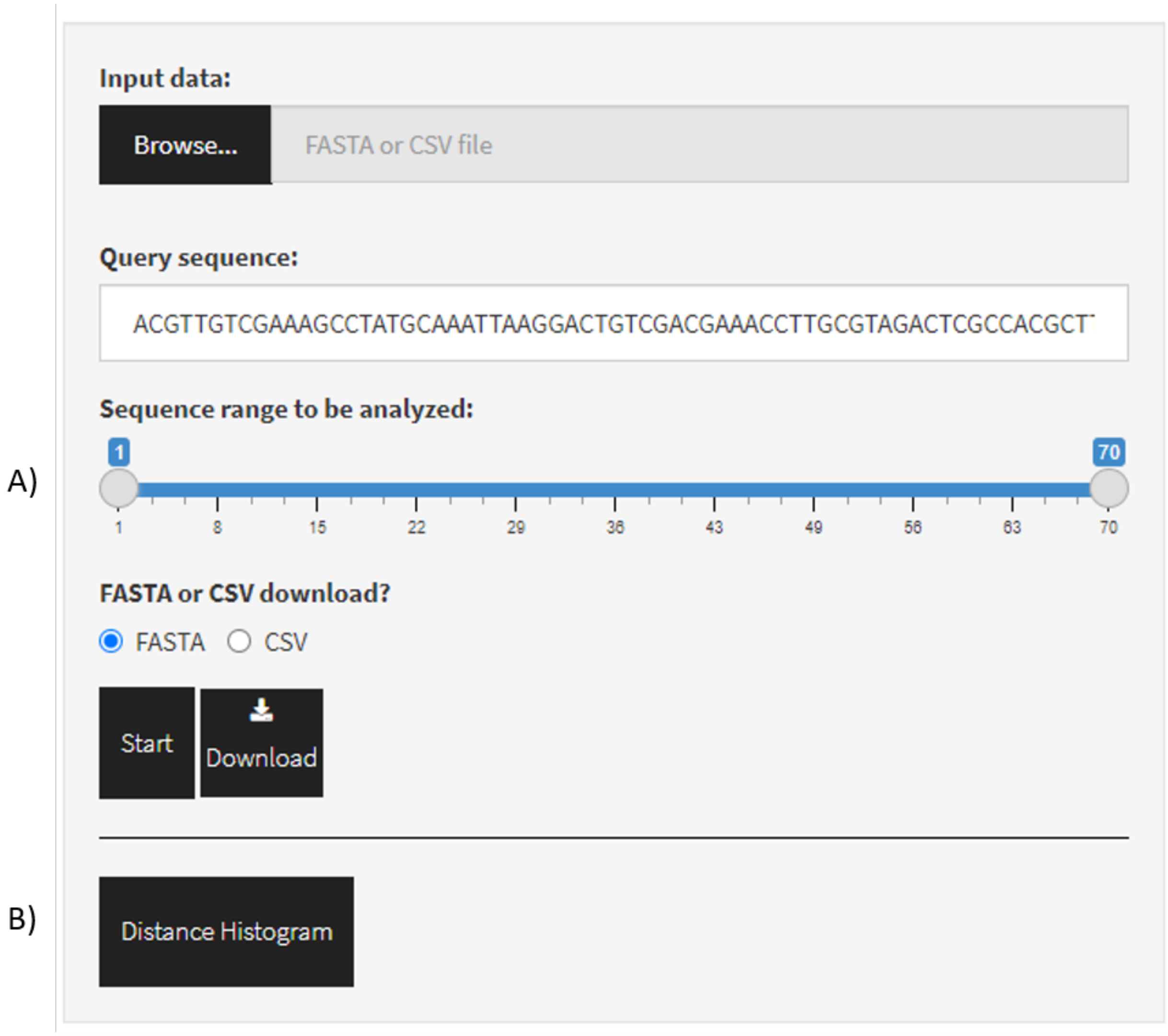

Figure 19: Screenshot of FASTAptameR-Distance.

The slider bar (**Fig. 19A**) allows the user to select a range of positions to query. For example, setting the two ends of this slider bar to 10 and 60 will truncate all of the sequences (**including the query sequence**) to be in that specific range. Thus, the resulting distance value will be the LED between positions 10-60 of the query sequence and positions 10-60 of every other sequence in the data. Thus, it is recommended that the starts of the sequences (5' ends, N-termini, *etc.*) are aligned.

A sample output data table is shown in **Fig. 20** with the following query sequence (the most abundant sequence from the 70HRT14 data set):

`ACGTTGTCGAAAGCCTATGCAAATTAAGGACTGTCGACGAAACCTTGCGTAGACTCGCCACGCTTGGTGT`.

Show 10 entries | Search:

| id | Rank | Reads | RPM | Distance | seqs |
|---|---|---|---|---|---|
| All | All | All | All | All | All |
| >1-417696-193358.44 | 1 | 417696 | 193358.44 | 0 | ACGTTGTCGAAAGCCTATGCAAATTAAGGACTGTCGACGAAACCTTGCGTAGACTCGCCACGCTTGGTGT |
| >5-74389-34435.91 | 5 | 74389 | 34435.91 | 1 | ACGTTGTCGAAAGCCTATGCAAATTAAGGACTGTCGACGAAACCTTGCGTAGACTCGCCGCGCTTGGTGT |
| >7-53608-24816.04 | 7 | 53608 | 24816.04 | 1 | ACGTTGTCGAAAGCCTATGCAAATTAAGGACTGTCGACGAAACCTTGCGTAGACTCGCCATGCTTGGTGT |
| >20-8003-3704.72 | 20 | 8003 | 3704.72 | 1 | ACGTTGTCGAAAGCCTATGCAAACTAAGGACTGTCGACGAAACCTTGCGTAGACTCGCCACGCTTGGTGT |
| >21-7815-3617.69 | 21 | 7815 | 3617.69 | 1 | ACGTTGTCGAAAGCCTATGCAAATCAAGGACTGTCGACGAAACCTTGCGTAGACTCGCCACGCTTGGTGT |
| >23-6177-2859.44 | 23 | 6177 | 2859.44 | 1 | ACGTTGTCGAAAGCCTATGCAGATTAAGGACTGTCGACGAAACCTTGCGTAGACTCGCCACGCTTGGTGT |
| >26-5487-2540.02 | 26 | 5487 | 2540.02 | 1 | ACGTTGTCGAAAGCCTATGCGAATTAAGGACTGTCGACGAAACCTTGCGTAGACTCGCCACGCTTGGTGT |
| >28-5302-2454.38 | 28 | 5302 | 2454.38 | 1 | ACGTTGTCGAAAGCCTGTGCAAATTAAGGACTGTCGACGAAACCTTGCGTAGACTCGCCACGCTTGGTGT |
| >29-5079-2351.15 | 29 | 5079 | 2351.15 | 1 | ACGTTGTCGAAAGCCTATGCAAATTGAGGACTGTCGACGAAACCTTGCGTAGACTCGCCACGCTTGGTGT |
| >31-4504-2084.98 | 31 | 4504 | 2084.98 | 1 | ACGTTGTCGAAAGCCTATGCAAATTAAGGACTGTCGACGAGACCTTGCGTAGACTCGCCACGCTTGGTGT |

Showing 1 to 10 of 72,921 entries | Previous 1 2 3 4 5 … 7,293 Next

Figure 20: FASTAptameR-Distance Output.

###### 3.7.3 Plotting

This module can also generate interactive histograms of distances (button shown in **Fig. 19B**, output shown in **Fig. 21**). The top plot corresponds to the distances between the query and *all unique sequences*, which allows greater visibility for low-abundance sequences. In contrast, the bottom plot corresponds to the distances between the query and all sequences, factoring in their read counts. In both cases, the query sequence is displayed at the top of the plot.

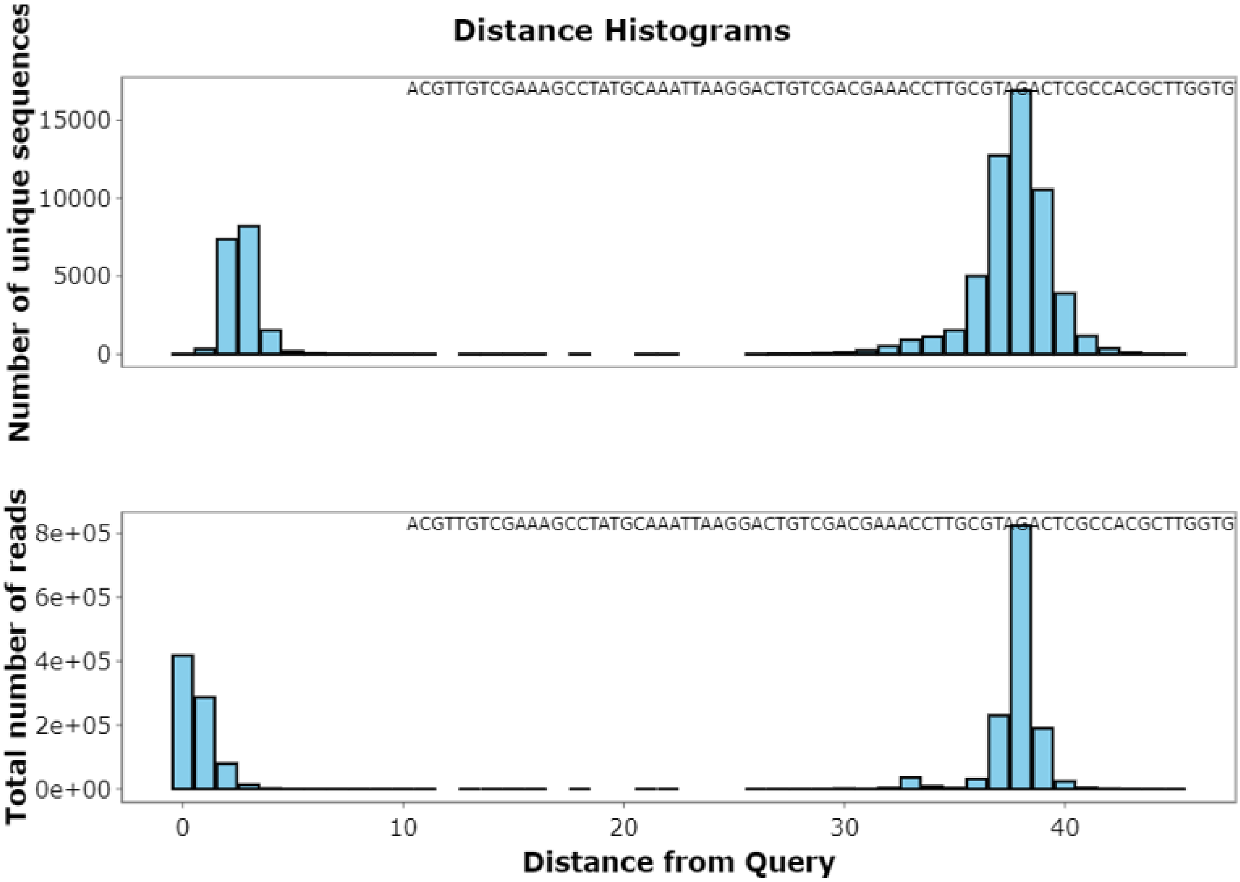

Figure 21: Distance Histograms with the 70HRT14 population as the target and the most abundant sequence as the query. A) Distance histogram with only unique sequences. B) Distance histogram with all reads. In both plots, the set of peaks on the left indicate the abundances of all sequences within a short edit distance of the seed sequence (that set constitutes a "cluster", see **section 3.10**), while the large gap between group and the rest of the population reflects the dissimilarities between any two molecules with 70 random positions.

#### 3.8 FASTAptameR-Enrich

##### 3.8.1 Description

The degree to which a given sequence enriches (or depletes) during a selection - or along different branches of a divergent evolution - is a powerful indicator of the relative fitness of that sequence. Enrichment analysis can be applied to the population as a whole or to subsets of closely related sequences (within-cluster).

FASTAptameR-Enrich calculates the enrichment (or depletion) of each sequence in one population relative to other populations. The module takes at least two *counted* FASTA files as input and returns a single data table after merging by sequences. Column headers for output data are appended with *.a*, *.b*, *.c*, *etc.*, depending on the order in which they are selected by the dropdown menu. Additional columns include enrichment scores (`E = RPM2 / RPM1`) and the base-two logarithm of the Enrichment ("log2(E)" = `log2(Enrichment)`). For simplicity, comparisons are only made between consecutive populations (*e.g.*, b:a, c:b, *etc.*). This output can be downloaded as a CSV.

All modules directly connected to FASTAptameR-Enrich are shown in **Fig. 22**, and a screenshot of the module interface is shown in **Fig. 23**.

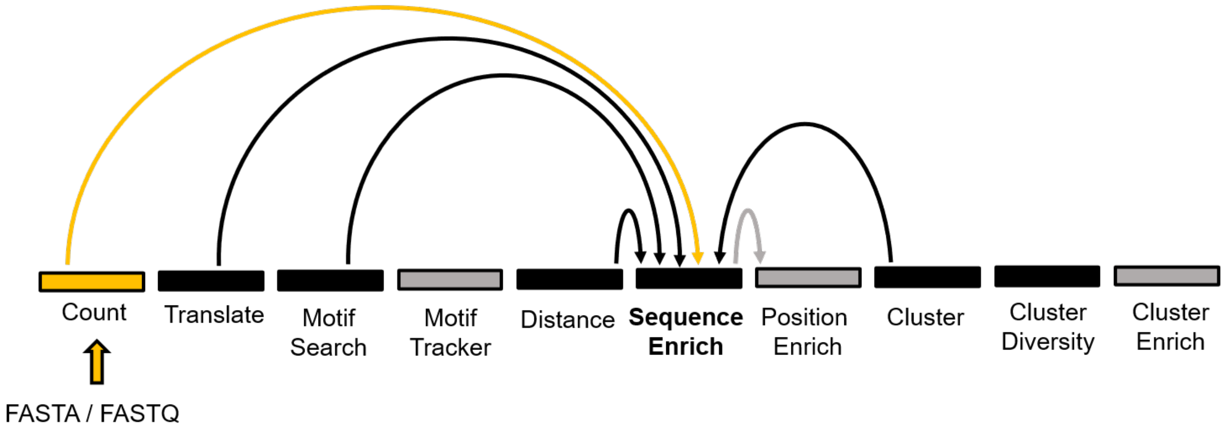

Figure 22: All modules connected to FASTAptameR-Enrich.

##### 3.8.2 Usage

The input FASTA files must be chosen with the file browser. The following set of radio buttons determine whether missing values are allowed in the output. Missing values result from sequences that are only present in a subset of the input files, such as when a sequence enriches from below the detection limit to above the detection limit.

The `Start` button begins the enrichment calculations, and the resulting data table will be shown on the right side of the screen. All numeric columns in this data table are filterable by typing into the corresponding text box (*e.g.*, `1 ... 10` to keep values in the range `[1:10]`) or by using the slider bar that is displayed after clicking in the corresponding text box. Note, these filters apply the mask only to the displayed data, so calculations will **not** be repeated when the filters are altered. To display all data again without the filters, simply delete the filters from the text boxes. Note, many other outputs are similarly filterable.

The `Download` button opens a file browser prior to downloading the output as a CSV file. A sample output data table that does not include missing sequences is shown in **Fig. 24**.

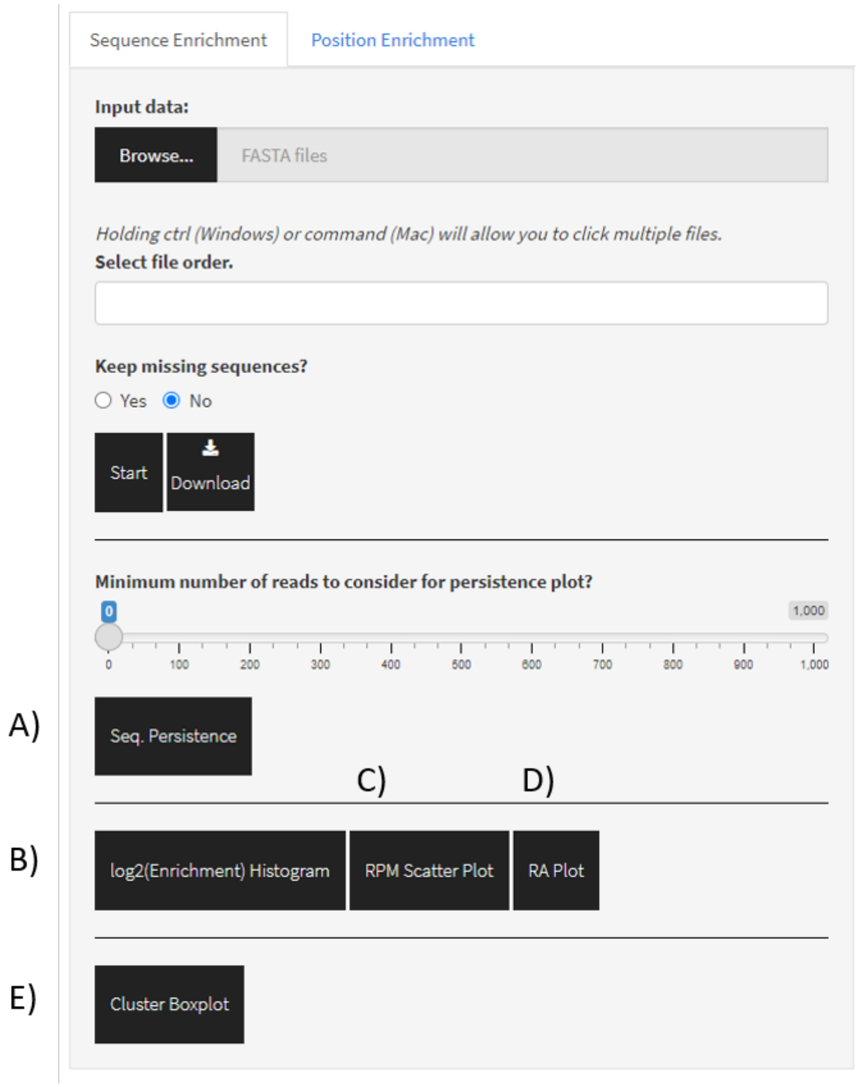

Figure 23: Screenshot of FASTAptameR-Enrich.

Show 10 entries Search:

| seqs | Rank.a | Reads.a | RPM.a | Rank.b | Reads.b | RPM.b | enrichment_ba |
|---|---|---|---|---|---|---|---|
| All | All | All | All | All | All | All | All |
| ACGTTGTCGAAAGCCTATGCAAATTAAGGACTGTCGACGAAACCTTGCGTAGACTCGCCACGCTTGGTGT | 1 | 417696 | 193358.44 | 3 | 161830 | 81408.87 | 0.421 |
| CATAGCGACTGTCCACGAATCCGAAGCCTAACGGGACAAAAGGCAAGAGCGCGATACCAATGCTGGACTG | 2 | 313312 | 145037.35 | 1 | 382391 | 192362.47 | 1.326 |
| AACCGCAAGCAACACCCAGCAAGAAACATCCGACGCACGACGGGAGAAAGTGCATTACCACGATGTCGAT | 3 | 174096 | 80591.94 | 5 | 104932 | 52786.23 | 0.655 |
| CATAGCGACTGCCCACGAATCCGAAGCCTAACGGGACAAAAGGCAAGAGCGCGATACCAATGCTGGACTG | 4 | 94978 | 43966.9 | 6 | 42954 | 21608.09 | 0.491 |
| ACGTTGTCGAAAGCCTATGCAAATTAAGGACTGTCGACGAAACCTTGCGTAGACTCGCCGCGCTTGGTGT | 5 | 74389 | 34435.91 | 9 | 32821 | 16510.66 | 0.479 |
| CCCTCCTTGTATGACGCTAACTGAGAATCCGAAGTCCAACGGGAGAAAGGACACTTATGACGTGGCGCG | 6 | 57625 | 26675.57 | 7 | 37701 | 18965.55 | 0.711 |
| ACGTTGTCGAAAGCCTATGCAAATTAAGGACTGTCGACGAAACCTTGCGTAGACTCGCCATGCTTGGTGT | 7 | 53608 | 24816.04 | 10 | 30749 | 15468.34 | 0.623 |
| AGCGCGGCACCCAAAATCGAAATCCGAAGGCGAACGGGAGAATGCGACCAAAGATACCCTGTGAATGGC | 8 | 39793 | 18420.84 | 15 | 15414 | 7754.04 | 0.421 |
| TTGACAATAACTCGAGAAGAACCGAGGTGCAAACGGGAGAACACAATGGATTACACCGAGCTCGGCTGAC | 9 | 33800 | 15646.58 | 4 | 136505 | 68669.08 | 4.389 |
| GCGAACCAAACCCAGATTACTAACCGTGGGCCTGAAACACGGGACAAAACAGGCATCAATGGAGTGGTAC | 10 | 29794 | 13792.14 | 176 | 732 | 368.23 | 0.027 |

Showing 1 to 10 of 21,924 entries Previous 1 2 3 4 5 … 2,193 Next

Figure 24: FASTAptameR-Enrich Output.

###### 3.8.3 Plotting

This module can generate five types of interactive plots (**Fig. 25, 26**): sequence persistence bar plots (button shown in **Fig. 23A**, output shown in **Fig. 25A**), $\log_2(Enrichment)$ histograms (one per comparison - button shown in **Fig. 23B**, output shown in **Fig. 25B**), RPM scatter plots (one per comparison - button shown in **Fig. 23C**, output shown in **Fig. 25C**), ratio average (RA) plots (one per comparison - button shown in **Fig. 23D**, output shown in **Fig. 25D**), and a cluster box plot in the case when **clustered** FASTA files are provided as input (button shown in **Fig. 23E**, output shown in **Fig. 26**).

The sequence persistence bar plot (**Fig. 25A**) bins sequences by the number of rounds in which they are found among all uploaded FASTA files. The slider bar just above this plot button filters these sequences by their respective read count. For the analysis shown, approximately 20000 sequences were found in both populations, whereas approximately 90000 sequences were found in only one population.

The spread of the $\log_2(Enrichment)$ histogram (**Fig. 25B**) relative to a vertical line at `x = 0` can indicate the overall magnitude of enrichment (or depletion), while displacement of the centroid of the distribution from this line can indicate possible directionality of the population's evolution. Similarly, the spread and displacement of the RPM scatter plot (**Fig. 25C**) with respect to the diagonal line at `y = x` can also indicate the magnitudes of enrichment and possible directionality. Finally, the RA plot (**Fig. 25D**) is used to show the relationship between the average log-RPM and $\log_2(Enrichment)$ for each sequence. Note that missing sequences (those with `RPM = 0` in one round) are treated as having `RPM = 0.1` for the sake of calculating their log2 values.

The cluster box plot (**Fig. 26**) shows the distribution of enrichment values for sequences after first grouping by cluster. The 25th and 75th quantiles are respectively represented by the bottom and top of each box. The line in the middle of the box represents the median. Whiskers are at most `1.5 * IQR`, and any points beyond that are shown as outliers. The red marker indicates where the seed sequence of the cluster falls. Individual points that are well above or well below the median represent species that are enriching or depleting relative to the cluster as a whole. Both types of outliers can be highly informative; species with strongly advantageous variations may be emerging as future dominant species for that cluster, while species with strongly disadvantageous variations can illuminate critical portions of the biomolecule.

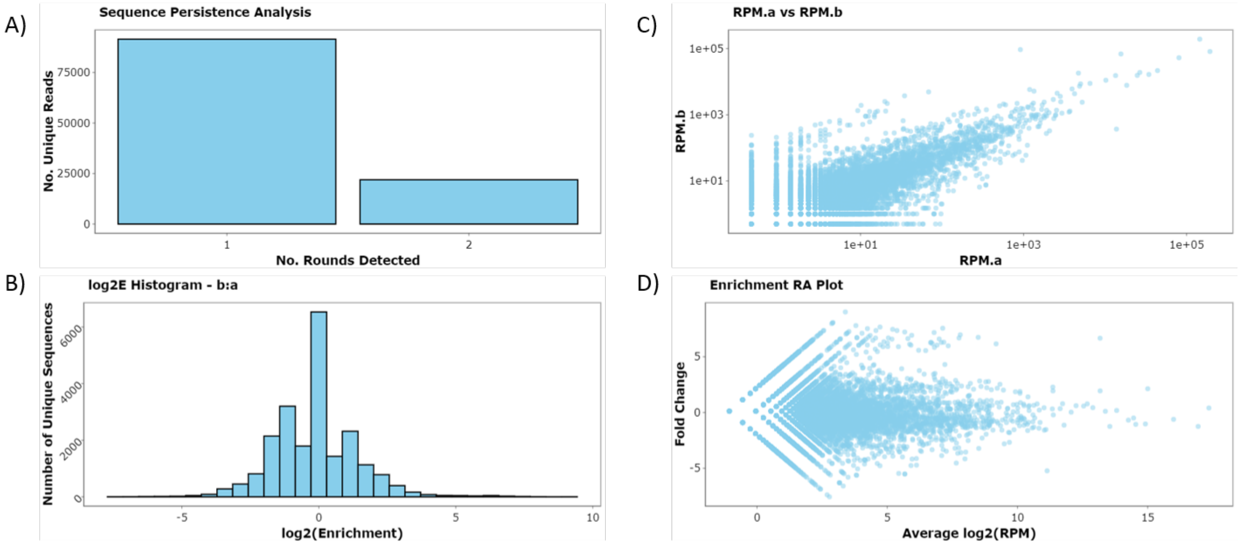

Figure 25: FASTAptameR-Enrich Plots. A) The sequence persistence bar plot shows how many rounds unique sequences persist. B) The histogram shows the distribution of fold-change between rounds. C) The RPM scatter plot shows the RPM of sequences across two rounds. D) The RA plot has fold change on the y-axis (`R` for Ratio) and average log2(RPM) on the x-axis (`A` for average). A small cloud of enriching sequences is visible in each of panel C (diagonal above the main distribution) and in panel D (horizontal above the main distribution).

#### 3.9 FASTAptameR-Position_Enrich

##### 3.9.1 Description

The plasticity of each position within a query sequence, as defined by its ability to tolerate substitutions, is related to the importance of that position and establishing the functional biomolecule. Sequences with mutations at critical positions are expected to deplete relative to those with mutations in more neutral locations. For aptamer selections, the functional core is often flanked by unimportant sequences that can be safely trimmed off once they are identified. However, identifying the functional boundaries experimentally can be labor intensive.

The FASTAptameR-Position_Enrich module calculates the average enrichment (or depletion) at each position for each species that does not match the corresponding residue in the user-defined reference sequence at that position. For example, if the first residue of the reference sequence is `E`, then this module will calculate the average enrichment of all sequences that do not have an `E` in the first position. Low 'enrichment' values imply low tolerance for substitution at those positions, which is typically interpreted as implying that those positions are important for function. Given the algorithm that computes these position-specific enrichment values, it is recommended that all sequences are of the same length (can be done by applying a filter to the output table from FASTAptameR-Count). Sequences with lengths different than the reference sequence will be omitted from these calculations.

This module accepts a CSV from the previous module (FASTAptameR-Enrich), though it exclusively operates on the `enrichment_ba` column. Thus, the output CSV file from an enrichment analysis of >2 populations can be uploaded here, but only the first enrichment column will be used.

The outputs of this module are two plots. The first is a bar plot showing the average enrichment values for each position. The second is a heat map that shows the average enrichment per position per residue.

All modules directly connected to FASTAptameR-Position_Enrich are shown in **Fig. 27**, and a screenshot of the module interface is shown in **Fig. 28**.

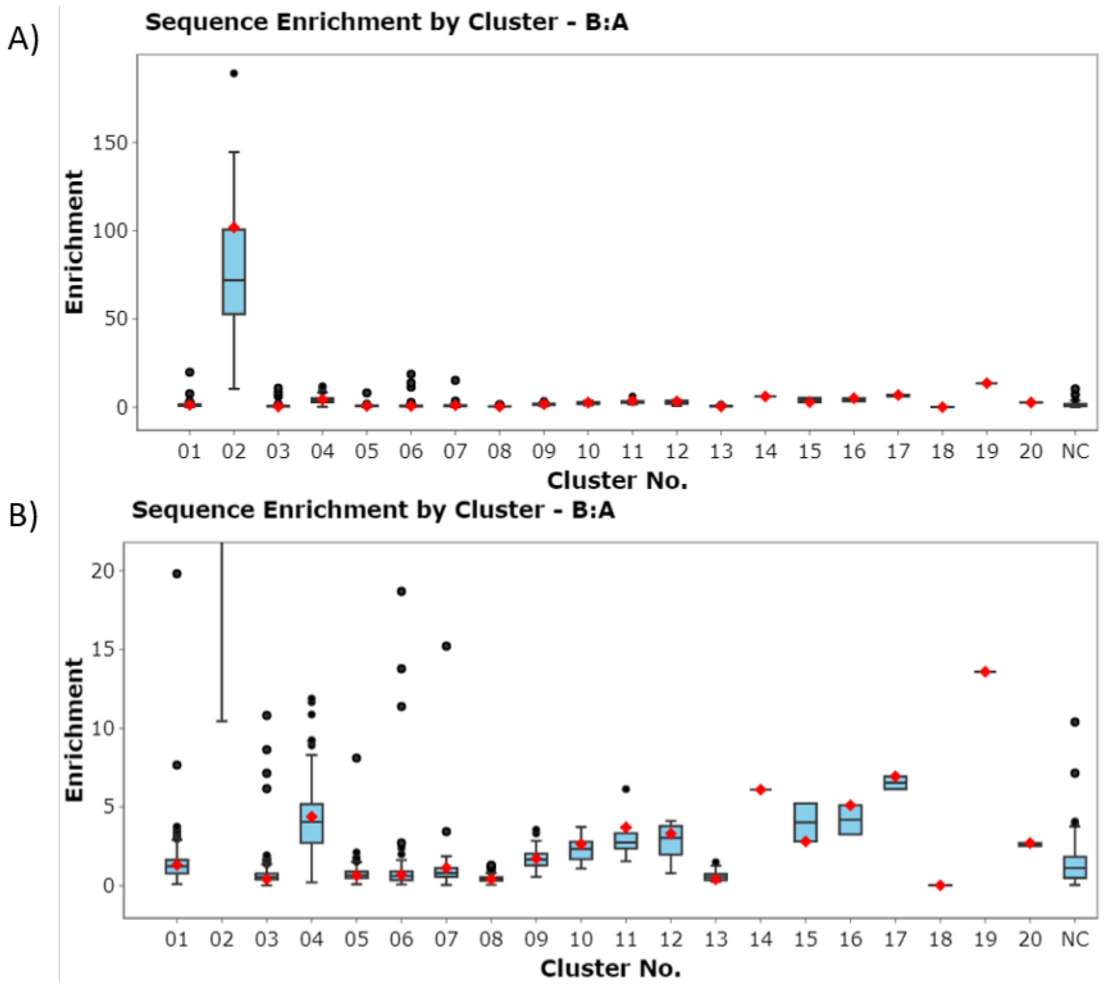

Figure 26: Cluster Box Plots of Sequence Enrichment. A) Enrichment distribution for the top 20 clusters. B) The same plot after zooming in on the y-axis.

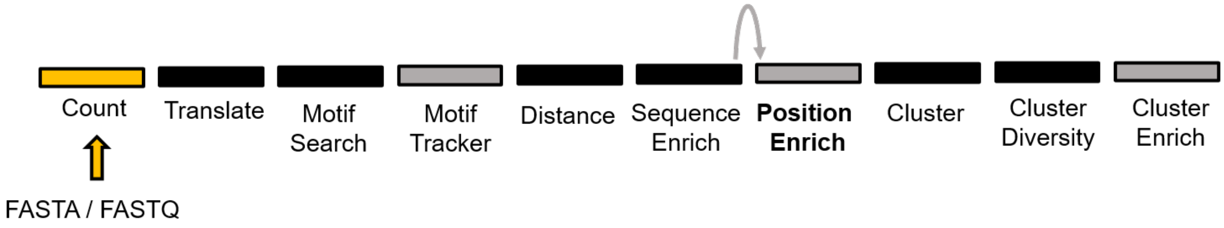

Figure 27: All modules connected to FASTAptameR-Position_Enrich.

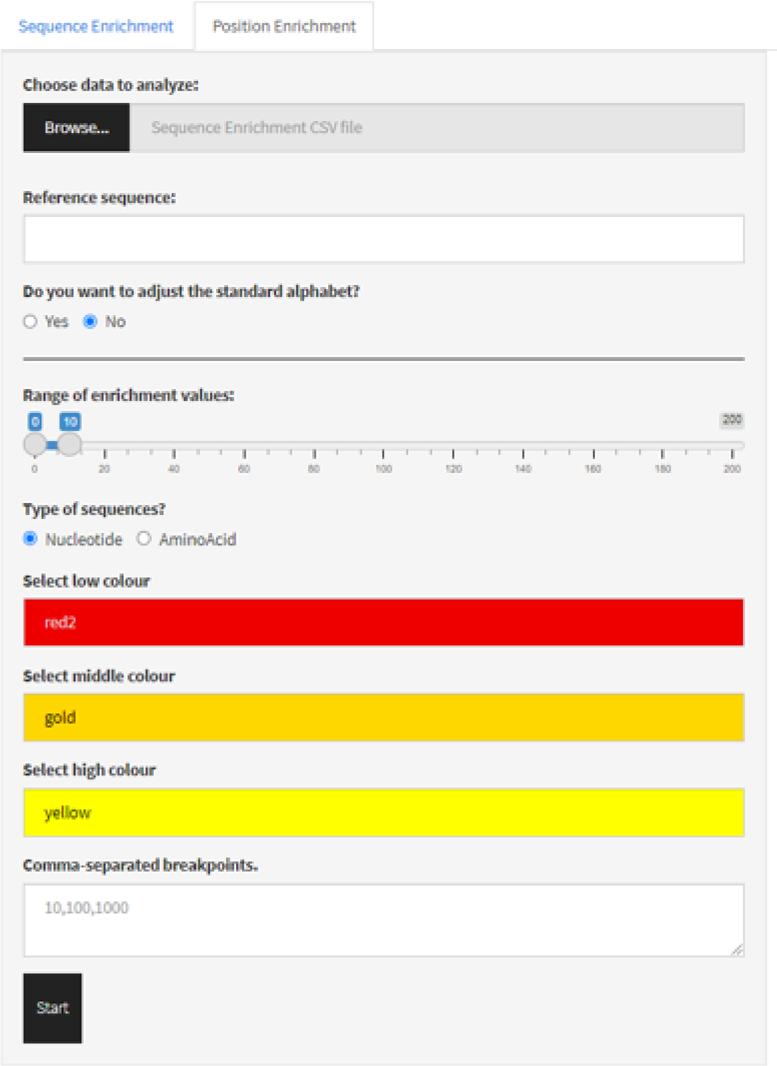

Figure 28: Screenshot of FASTAptameR-Position_Enrich. Note that the three color choices are used to create a gradient, and the midpoint is set at the 65th quantile.

##### 3.9.2 Usage

The input CSV file must be chosen with the file browser, and the reference sequence must be added in the subsequent text box.

The first set of radio buttons allows the standard alphabet to be altered to include nonstandard nucleotides or amino acids. Each change should occupy a single line. To add a residue, enter its single-letter code. To replace a residue, enter a comma-separated pair (*e.g.*, `U,F` will replace `U` with `F` in the algorithm and resulting plots).

The slider bar allows the user to set the minimum and maximum enrichment values (*e.g.*, `0-10` means that any value greater than 10 is made equal to 10 for the plot). The final set of radio buttons determines whether the algorithm searches for nucleotide or amino acid residues. The next three text boxes allow the user to set the "low", "middle", and "high" colors for the plots.

The text input at the bottom of the UI allows the user to enter comma-separated breakpoints for which average enrichments will be calculated. For example, if the user enters `1,20,50` and the reference sequence is 70 nucleotides, then position-specific enrichments will be averaged for positions [1,20), [20, 50), and [50, 70).

Finally, the `Start` button generates the two plots.

##### 3.9.3 Plotting

FASTAptameR-Position_Enrich generates two types of plots (**Fig. 29**): 1) average enrichment bar plot and 2) average enrichment heat map. When the mouse hovers over a bar in either plot, the position and enrichment value is returned. The bar plot (**Fig. 29A**) shows the reference sequence on the x-axis and the average enrichment on the y-axis. The heat map (**Fig. 29B**) shows the reference sequence on the x-axis, possible residues on the y-axis, and average enrichment in the color axis.

The motif of interest for this example is the family 1 pseudoknot (F1Pk), which is defined as `UCCG[n*]CGGGAnAAAA` where `n` is any nucleotide and `[n*]` is any number (typically 4-10) of any nucleotide (C. Tuerk, Macdougal, and Gold 1992; Ditzler MA 2013). The F1Pk is shown between the dashed lines in **Figs. 29A,B**. **Fig. 29C** shows the experimentally determined secondary structure of this motif. The workflow to generate these plots is given below:

1. Count 70HRT14.fastq and 70HRT15.fastq with FASTAptameR-Count.
2. Generate top five clusters for each counted population with FASTAptameR-Cluster. Include all sequences (min. reads > 0) and use LED = 7.
3. Use cluster 2 from 70HRT14 and cluster 1 from 70HRT15 in FASTAptameR-Enrich. Representative sequences from each population contain the F1Pk motif.
4. Use the output CSV in FASTAptameR-Position_Enrich, specifying the most abundant sequence in 70HRT15 as the reference sequence.

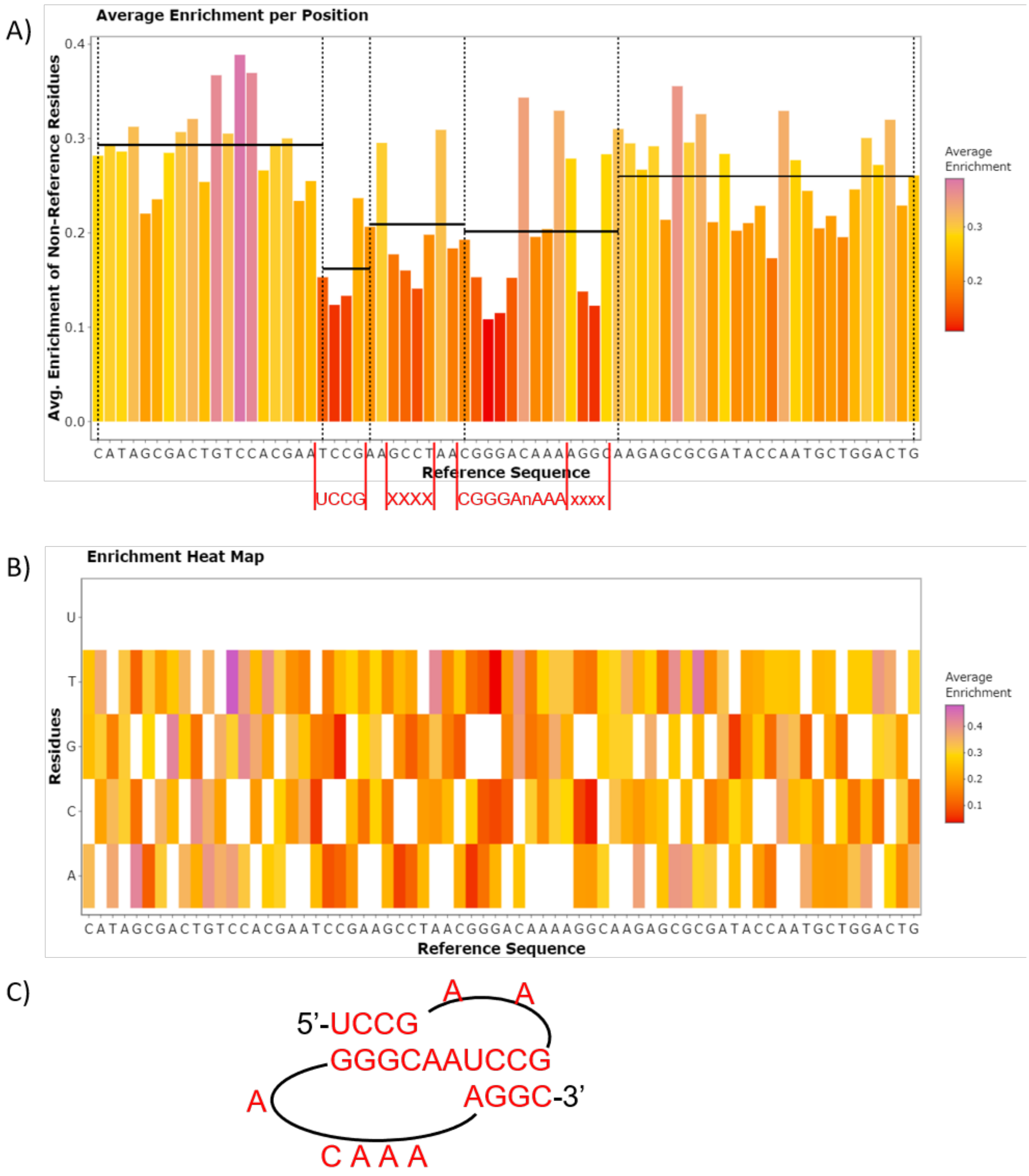

Figure 29: FASTAptameR-Position_Enrich Plots. A) The heat map shows the user-defined reference sequence on the x-axis and all possible residues on the y-axis (nucleotides for this use case). Colors depict average enrichment of non-reference residues. B) The bar plot shows the user-defined reference sequence on the x-axis and average enrichment of non-reference residues on the y-axis. Horizontal dashed lines indicate average enrichment values (left-inclusive) within the ranges of positions 1-19, 20-42, and 43-70. Positions ~20-43 have low enrichment scores, suggesting the importance of nucleotides within this region, which contains the F1Pk motif. C) The experimentally determined structure the F1Pk module within the most abundant sequence of 70HRT15.

##### 3.10 FASTAptameR-Cluster

###### 3.10.1 Description

Combinatorial selections often produce clusters of very closely related sequences as a result of divergent evolution (accumulation of point mutations, neutral drift), or convergent evolution (independent seeding of the population with closely related sequences, as with low-level mutatgenesis of a seed sequence or dense oversampling at a limited number of positions). Grouping sequences into clusters can greatly simplify analysis of population structure and evolution outcomes/dynamics. Sequence clusters often behave as quasi-species, sampling local sequence space and evolving in similar fashion in response to local fitness landscapes. The FASTAptameR-Cluster module defines sequence clusters within the population, while the other modules in this platform - FASTAptameR-Cluster_Diversity, FASTAptameR-Cluster_Enrich, and the box plot feature of FASTAptameR-Enrich - allow the user to explore different aspects of the quasi-species nature of the clusters.

FASTAptameR-Cluster groups sequences according to sequence relatedness for all sequences in the population within a user-defined threshold of similarity. The module accepts a *counted* FASTA file as input. If no output directory is specified (the default setting), the module returns a *clustered* data table to the screen. This data table contains all sequences and clusters and can be downloaded as a single FASTA or CSV file. If an output directory is specified, then no data table will be created, and one FASTA file per cluster will be written to the output directory, up to a user-defined number of output files.

Briefly, the module identifies clusters in an iterative manner. During each iteration, the most abundant sequence that has not yet been clustered becomes a cluster "seed" for that iteration. Any other sequences that have not yet been clustered and that are within a user-defined edit distance of this seed sequence are added to this cluster. This process repeats until all sequences are clustered or a predefined number of clusters is created.

All modules directly connected to FASTAptameR-Cluster are shown in **Fig. 30**, and a screenshot of the module interface is shown in **Fig. 31**.

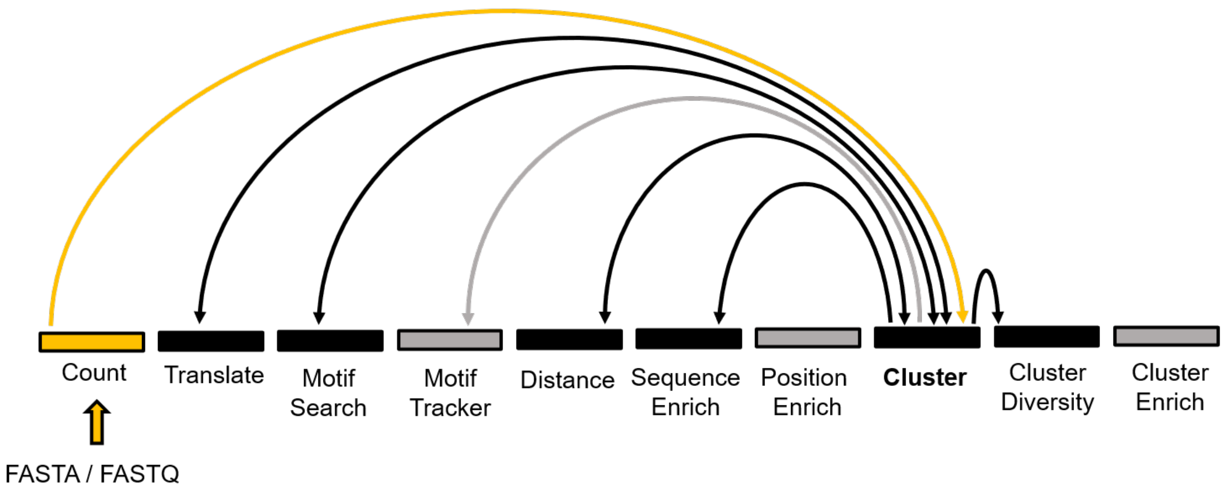

Figure 30: All modules connected to FASTAptameR-Cluster.

###### 3.10.2 Usage

The input FASTA file must be chosen with the file browser. The first slider bar (**Fig. 31A**) sets the minimum number of reads a sequence must have for it to be included within a cluster (`DEFAULT = 10`). Sequences with the chosen number or fewer reads are removed prior to clustering. Thus, setting this filter

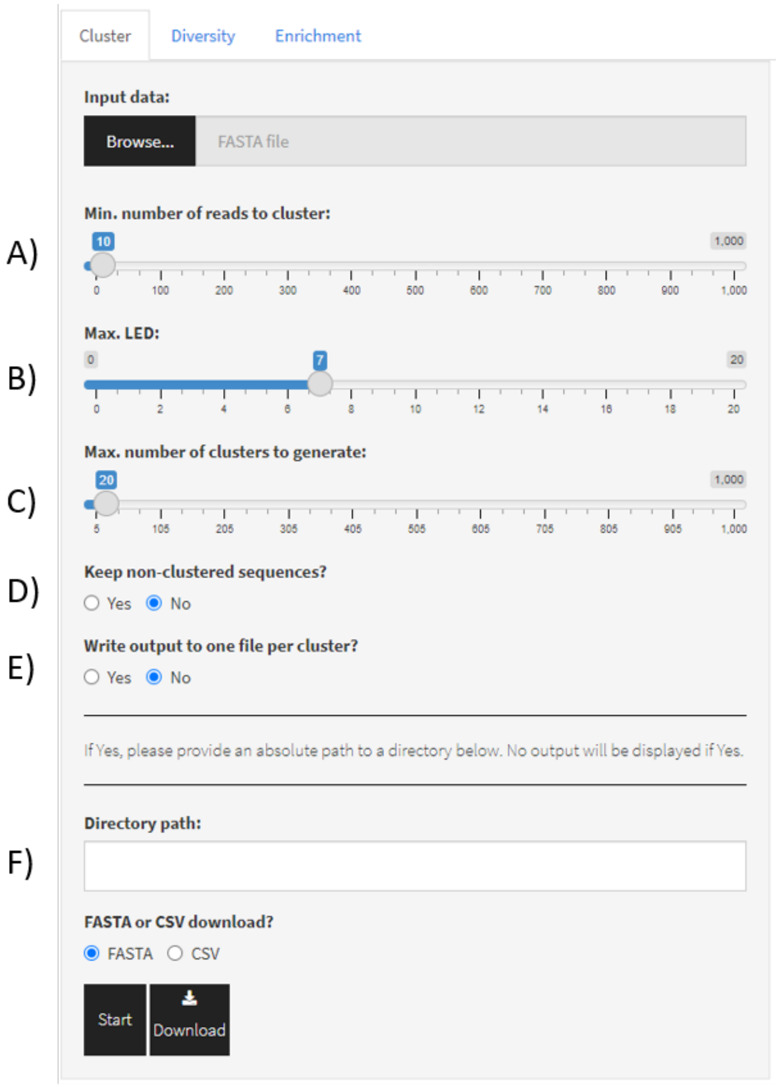

Figure 31: Screenshot of FASTAptameR-Cluster.

to values >1 can significantly shorten runtime, especially for relatively complex data sets. The second slider bar (**Fig. 31B**) sets the maximum Levenshtein edit distance to consider between a seed sequence and all other sequences (`DEFAULT = 7`). Users may wish to run FASTAptameR-Distance to guide threshold LED values to use in establishing cluster definitions (see **Fig. 21**) The third slider bar (**Fig. 31C**) sets the total number of desired clusters (`DEFAULT = 20` to limit runtime during exploratory runs). Note, any remaining sequences will be grouped as `NC` ("not clustered").

The first set of radio buttons (**Fig. 31D**) indicates whether non-clustered sequences should be kept (`DEFAULT = No`). If `Yes` then the sequence IDs of non-clustered sequences will be appended with `NC`.

The second set of radio buttons (**Fig. 31E**) indicate whether each cluster should be written to a different FASTA file (`DEFAULT = No`). If `No`, then all clusters are grouped together and can be and downloaded in a single file. If `Yes`, then each cluster will be written to its own FASTA file, and no data table will be displayed. Note that if this option is `Yes`, then a directory path **must be copied or typed** into the corresponding text box (**Fig. 31F**) if this option is `Yes`. A sample directory path could be `C:/Users/Kramer/Desktop/Data/directory/`, though this will depend on your system. **Please note** that this requires backslashes (`/`) and that forward slashes (`\`) will cause errors. Also note that the path should end with a backslash.

The `Start` button will begin the clustering process. The results will be displayed as a data table on the right side of the screen. The `Download` button opens a file browser prior to downloading the output as a FASTA or CSV file (`DEFAULT = FASTA`, which is required for subsequent modules).

Algorithm progress will be shown below these buttons and will update after each cluster finishes. These notifications occur regardless of whether the module is writing to one or many files.

A sample output data table is shown in **Fig. 32**.

Show 10 entries Search:

| id | Rank | Reads | RPM | cluster | rankInCluster | LED | seqs |
|---|---|---|---|---|---|---|---|
| All | All | All | All | All | All | All | All |
| >1-417696-193358.44-1-1-0 | 1 | 417696 | 193358.44 | 1 | 1 | 0 | ACGTTGTCGAAAGCCTATGCAAATTAAGGACTGTCGACGAAACCTTGCGTAGACTCGCCACGCTTGGTGT |
| >2-313312-145037.35-2-1-0 | 2 | 313312 | 145037.35 | 2 | 1 | 0 | CATAGCGACTGTCCACGAATCCGAAGCCTAACGGGACAAAAGGCAAGAGCGCGATACCAATGCTGGACTG |
| >3-174096-80591.94-3-1-0 | 3 | 174096 | 80591.94 | 3 | 1 | 0 | AACCGCAAGCAACACCCAGCAAGAAACATCCGACGCACGACGGGAGAAAGTGCATTACCACGATGTCGAT |
| >4-94978-43966.9-2-2-1 | 4 | 94978 | 43966.9 | 2 | 2 | 1 | CATAGCGACTGCCCACGAATCCGAAGCCTAACGGGACAAAAGGCAAGAGCGCGATACCAATGCTGGACTG |
| >5-74389-34435.91-1-2-1 | 5 | 74389 | 34435.91 | 1 | 2 | 1 | ACGTTGTCGAAAGCCTATGCAAATTAAGGACTGTCGACGAAACCTTGCGTAGACTCGCCGCGCTTGGTGT |

Figure 32: FASTAptameR-Cluster Output.

Note that the new *id* column is the old *id* with three new values: `Cluster Number`, `Rank in Cluster`, and `Distance to Cluster Seed` - appended onto the original three identifiers.

#### 3.11 FASTAptameR-Cluster_Diversity

##### 3.11.1 Description

FASTAptameR-Cluster_Diversity evaluates diversity across the *clustered* population and sequence relationships within and between clusters. The module accepts a *clustered* FASTA file as input and returns a data table with metadata for each cluster. This data table can be downloaded as a CSV file.

All modules directly connected to FASTAptameR-Cluster_Diversity are shown in **Fig. 33**, and a screenshot of the module interface is shown in **Fig. 34**.

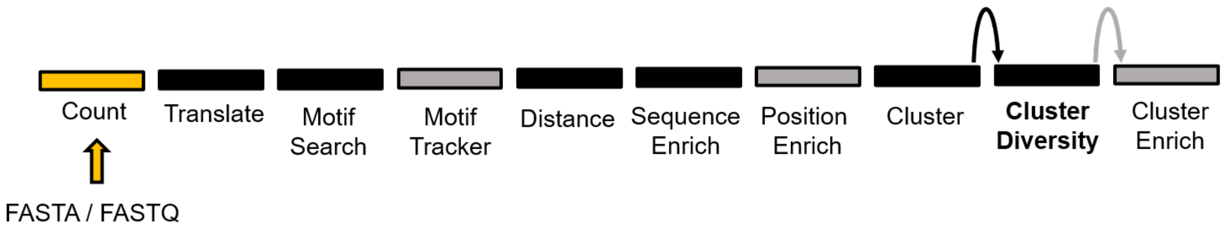

Figure 33: All modules connected to FASTAptameR-Cluster_Diversity.

##### 3.11.2 Usage

The input FASTA file (clustered FASTA file from FASTAptameR-Cluster) must be chosen with the file browser. The `Start` button begins the analysis. The results will be displayed as a data table on the right side of the screen and will include the following columns: `Cluster Number`, `Seed Sequence`, `Total Sequences`, `Total Reads`, and `Total RPM`. The `Download` button opens a file browser prior to downloading the output as a CSV file, which can be used by FASTAptameR-Cluster_Enrich. A sample output data table is shown in **Fig. 35**.

##### 3.11.3 Plotting

This module can generate metaplots of the analyzed data. These line plots correspond to the number of unique sequences per cluster, total reads per cluster, and average LED to seed sequence per cluster (button shown in **Fig. 34A**, output plots shown in **Fig. 36A**).

This module is also able to analyze clusters by converting all sequences into k-mer vectors (see below) and rendering an interactive 2D PCA plot, colored by cluster (button shown in **Fig. 34B**, output shown in **Fig. 36B**). The value of `k` can be chosen with the first set of radio buttons (`DEFAULT = 3`). The slider bar indicates how many of the top clusters should be plotted (max = 21 clusters due to graphics limitations). The second set of radio buttons indicates whether non-clustered (`NC`) sequences should be plotted (`DEFAULT = Yes`). Note that non-clustered (`NC`) sequences in the output are marked as `NA` in this plot.

Only nucleotide sequences without ambiguities should be plotted in this module. The large k-mer matrix needed for peptide sequences may return errors related to memory usage. Further, this module will alter any set of sequences with characters outside of `[A, C, G, T/U]` by converting any other character to `X`.

##### 3.11.4 A side note on how k-mer clustering works and what it means

In the k-mer method, sequence relatedness is measure by the degree to which sequences share a set of short sequence elements ('k-mers'), typically of length 3, 4, or 5. For example, if k=3, there are 64 possible k-mers: 'AAA', 'AAC', 'AAG', 'AAT', 'ACA', 'ACC', . . . , 'TTA', 'TTC', 'TTG', 'TTT.' *These triplet sequences constitute the axes of a 64-dimensional hyperspace.*

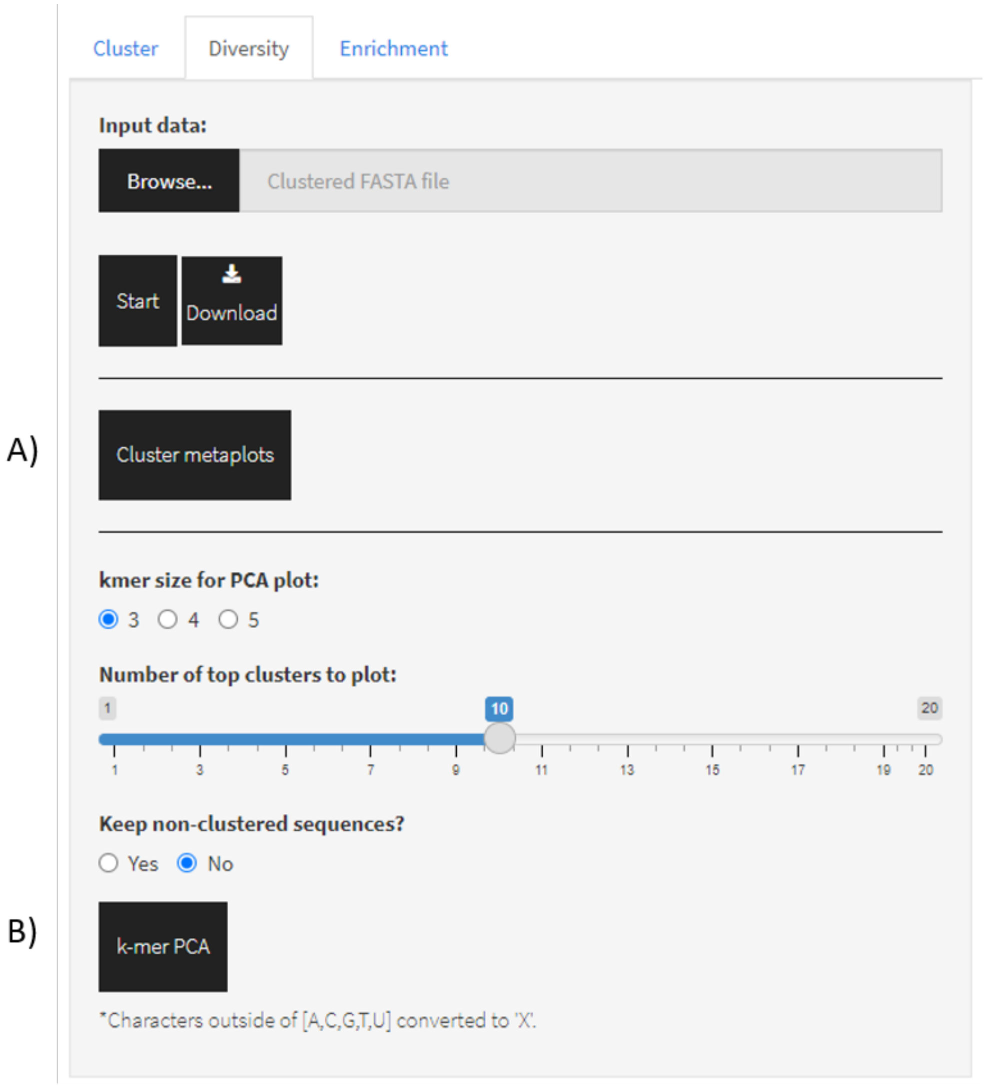

Figure 34: Screenshot of FASTAptameR-Cluster_Diversity.

Show 10 entries  Search:

| Cluster | Seeds | TotalSequences | TotalReads | TotalRPM | AverageLED |
|---|---|---|---|---|---|
| All | All | All | All | All | All |
| 1 | ACGTTGTCGAAAGCCTATGCAAATTAAGGACTGTCGACGAAACCTTGCGTAGACTCGCCACGCTTGGTGT | 1259 | 770383 | 356623.54 | 1.86 |
| 2 | CATAGCGACTGTCCACGAATCCGAAGCCTAACGGGACAAAAGGCAAGAGCGCGATACCAATGCTGGACTG | 1295 | 652468 | 302038.75 | 1.97 |
| 3 | AACCGCAAGCAACACCCAGCAAGAAACATCCGACGCACGACGGGAGAAAGTGCATTACCACGATGTCGAT | 528 | 257675 | 119282.23 | 1.63 |
| 4 | CCCTCCTTGTATGACGCTAACTGAGAATCCGAAGTCCAACGGGAGAAAGGACACTTATGACGTGGCGCG | 324 | 101448 | 46962.14 | 1.49 |
| 5 | AGCGCGGCACCCAAAATCGAAATCCGAAGGCGAACGGGAGAATGCGACCAAAGATACCCTGTGAATGGC | 262 | 73809 | 34167.47 | 1.47 |
| 6 | TTGACAATAACTCGAGAAGAACCGAGGTGCAAACGGGAGAACACAATGGATTACACCGAGCTCGGCTGAC | 275 | 67908 | 31435.87 | 1.51 |
| 7 | GCGAACCAAACCCAGATTACTAACCGTGGGCCTGAAACACGGGACAAAACAGGCATCAATGGAGTGGTAC | 148 | 41566 | 19241.67 | 1.16 |
| 8 | ACGTTGTGCACGGATGCCCACGGTCGCACGAAACCTTGTGTGGGATAGCGCGAATACTGACGAGTGTGCC | 163 | 44693 | 20689.16 | 1.26 |
| 9 | ACCAAATCCCGAACTACAAATCCGAACGCTAACGGGACAATTGCGAAATGGAACATACGGGCCTGGTTGA | 67 | 9114 | 4219.07 | 1.1 |
| 10 | GTGCGCTACCACATGATCCGAGGCAAAACGGGAAAAGATAGCATCGATTACGGAACCGGCCACGCACA | 54 | 7292 | 3375.58 | 0.98 |

Showing 1 to 10 of 21 entries  Previous 1 2 3 Next

Figure 35: FASTAptameR-Cluster_Diversity Output.

For each sequence, count the number of occurrences for each k-mer. *These counts constitute the values associated with the axes for that sequence.* For example, the sequence 'AAAAGT' has 2 copies of 'AAA', 1 copy of 'AAG', 1 copy of 'AGT', and 0 copies of all other triplets. Every sequence can therefore be defined as a single point in 64-dimensional space, with the (x, y, z, . . . ) values corresponding to the respective k-mer counts.

More generally, every sequence can therefore be defined as a single point (or 'vector') in $A^k$**-dimensional space**, where **A** is the number of letters in the polymer alphabet [standard values = 4 for nucleic acids and 20 for proteins, but other values are also possible], and $k$ is the chosen k-mer value.

The DISTANCE between two sequences is then simply the Euclidean distance calculated from one point to another in that hyperdimensional space.

Don't get thrown off by the word 'hyperdimensional'. It's really easy to see the pattern and then generalize from there. Consider going from 2D to 3D. The distance between two points in 2D is just the Pythagorean formula that you've known for many years:

$$d = \sqrt{([x_2 - x_1]^2 + [y_2 - y_1]^2)}.$$

Applying this to 3D, you simply add the z-dimension:

$$d = \sqrt{([x_2 - x_1]^2 + [y_2 - y_1]^2 + [z_2 - z_1]^2)}.$$

As you add more dimensions, you just add more variables.

But what do these distances mean?'

The biologically inclined among us are used to thinking of mutational or evolutionary distance, which is essentially the number of mutational events that are thought to have occurred since the two species diverged. k-mer distance is not mutational distance, but the two kinds of 'distance' are related, in that the k-mer analysis gives longer distances for more divergent sequences, so long as "k" is large enough to provide the necessary resolution.

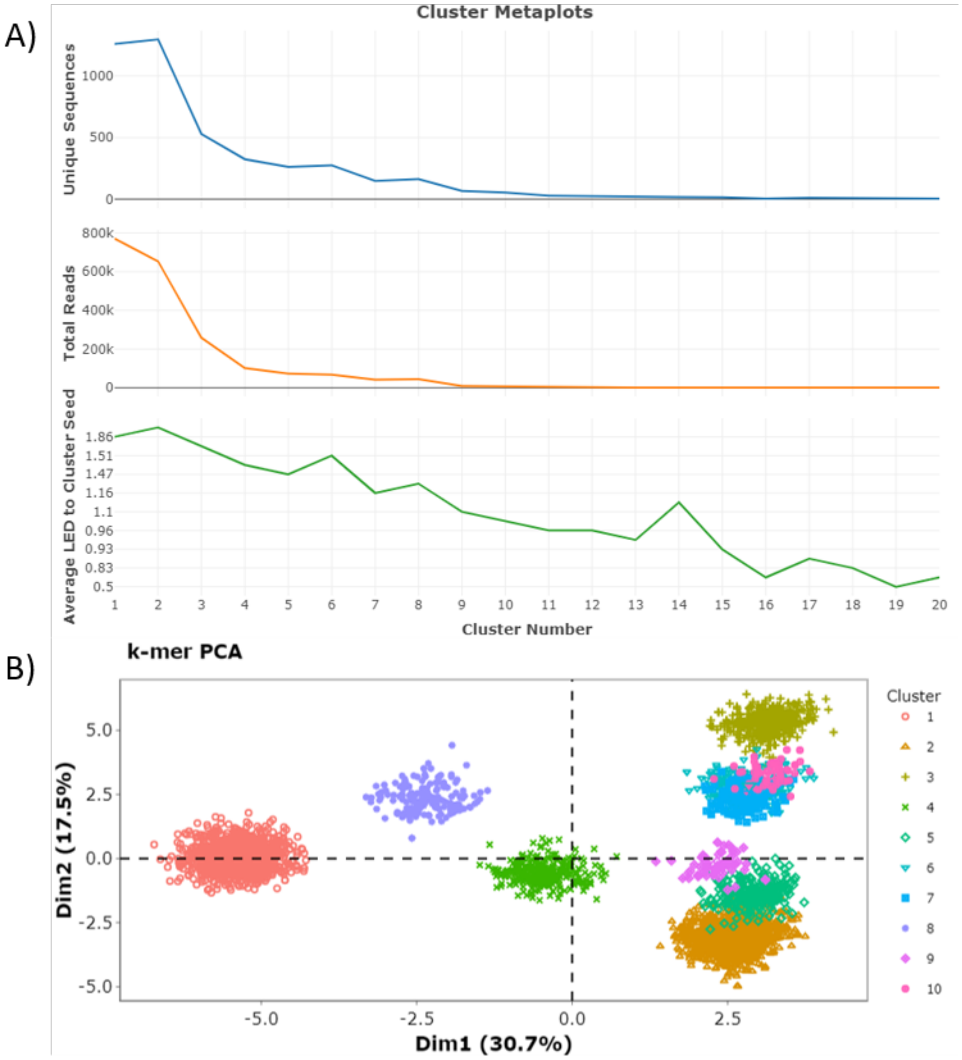

Figure 36: Cluster metaplots. A) Cluster metaplots depict number of unique sequences, total number of reads, and average LED to cluster seed per cluster. B) The k-mer PCA plot can qualitatively suggest how well the cluster algorithm performed (as indicated by cohesive grouping of each cluster within the plot) and identifies clusters that are especially distinct from (well-separated) or similar to (close or overlapping) the others.

#### 3.12 FASTAptameR-Cluster_Enrich

##### 3.12.1 Description

Considering all the members of a cluster together (rather than as independently evolving species) can help the user spot large-scale trends, in addition to adding statistical weight to enrichment analyses. For example, a cluster with a large number of low-abundance functional variants may outperform another cluster with few variants that are each at higher abundances. The strong performance of the first cluster might be missed in an analysis that looks only at individual species. A counterpoint to this collective approach is provided in the cluster box plot feature of FASTAptameR-Enrich (see **Fig. 26**).

FASTAptameR-Cluster_Enrich calculates the enrichment (or depletion) of each cluster in one population relative to other populations. Thus, this module is conceptually identical to FASTAptameR-Enrich but is applied to clusters rather than individual sequences. The module accepts two or three *cluster-analysis* CSV files as input and returns a data table after merging by `Seed Sequence`. Thus, this module assumes that cluster seeds are consistent across populations, though this may not always be a valid assumption.

The output data tables can be downloaded as CSV files. The first CSV file provides summary statistics for each cluster in each population. The second CSV file contains enrichment scores.

All modules directly connected to FASTAptameR-Cluster_Enrich are shown in **Fig. 37**, and a screenshot of the module interface is shown in **Fig. 38**.

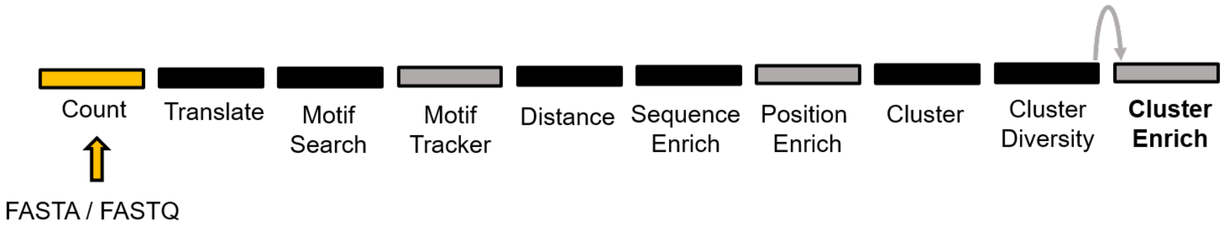

Figure 37: All modules connected to FASTAptameR-Cluster_Enrich.

##### 3.12.2 Usage

The input CSV files must be chosen with the file browser. The `Start` button begins the enrichment calculation. The results will be displayed as two data tables on the right side of the screen. The first table summarizes each cluster, and the second table provides enrichment values. The `Download` button opens a file browser prior to downloading the output as a CSV file.

A sample output data table is shown in **Fig. 39**.

Note that columns 3-5 and 7-9 refer to *total* values in the given cluster.

##### 3.12.3 Plotting

After merging by seed sequence, this module will generate a line plot in which the x-axis corresponds to population, and the y-axis corresponds to the total RPM of the seed's cluster (**Fig. 40**). Although only two populations were used here for illustration, this tool can be especially useful for observing the rise and fall of multiple clusters over the course of multiple rounds of selection.

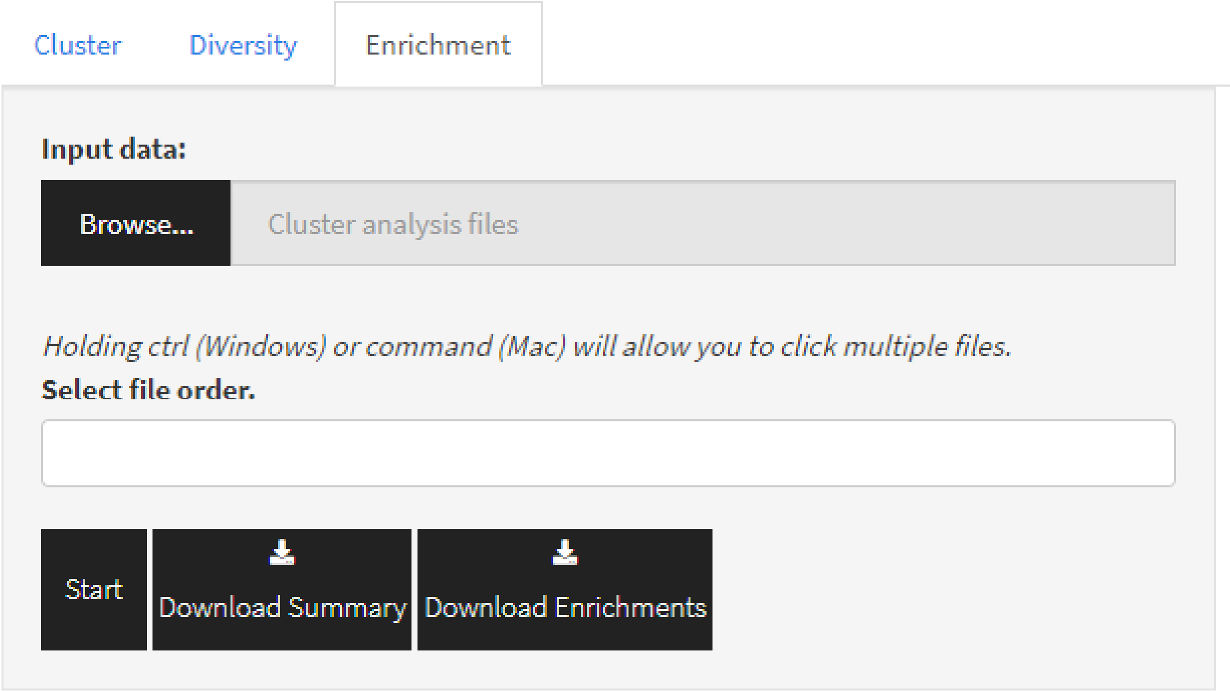

Figure 38: Screenshot of FASTAptameR-Cluster_Enrich.

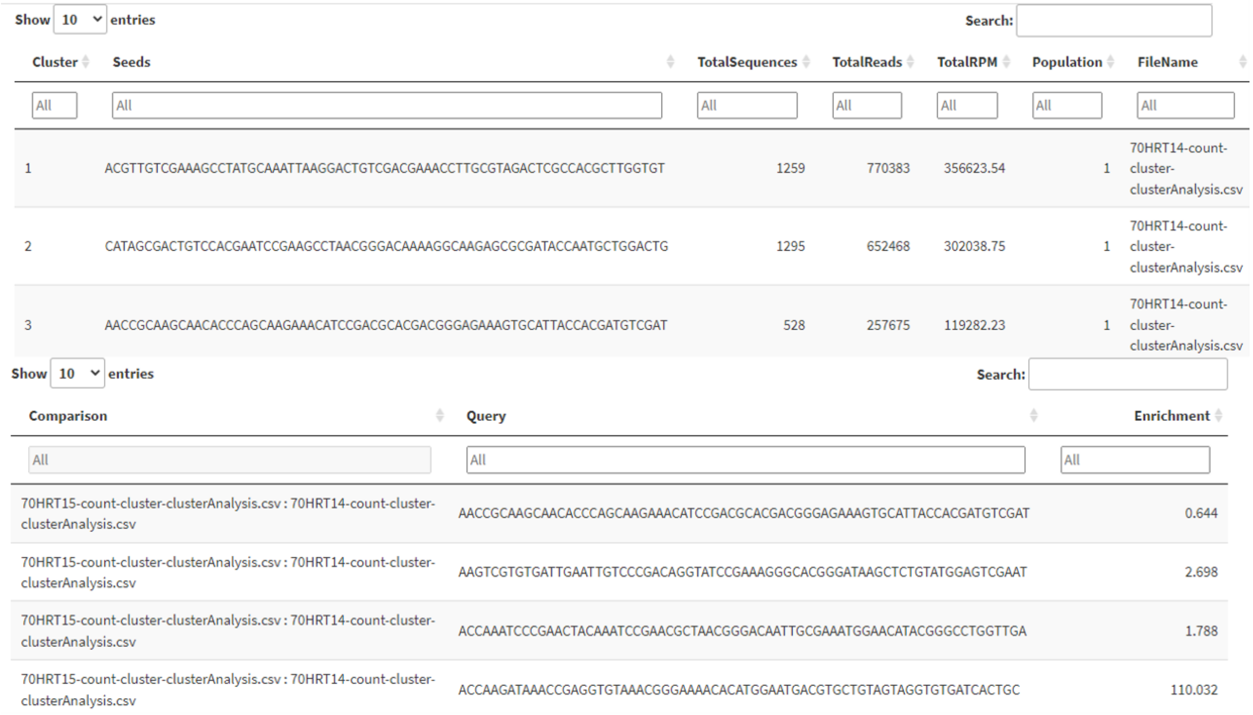

| Cluster | Seeds | TotalSequences | TotalReads | TotalRPM | Population | FileName |
|---|---|---|---|---|---|---|
| All | All | All | All | All | All | All |
| 1 | ACGTTGTCGAAAGCCTATGCAAATTAAGGACTGTCGACGAAACCTTGCGTAGACTCGCCACGCTTGGTGT | 1259 | 770383 | 356623.54 | 1 | 70HRT14-count-cluster-clusterAnalysis.csv |
| 2 | CATAGCGACTGTCCACGAATCCGAAGCCTAACGGGACAAAAGGCAAGAGCGCGATACCAATGCTGGACTG | 1295 | 652468 | 302038.75 | 1 | 70HRT14-count-cluster-clusterAnalysis.csv |
| 3 | AACCGCAAGCAACACCCAGCAAGAAACATCCGACGCACGACGGGAGAAAGTGCATTACCACGATGTCGAT | 528 | 257675 | 119282.23 | 1 | 70HRT14-count-cluster-clusterAnalysis.csv |

| Comparison | Query | Enrichment |
|---|---|---|
| All | All | All |
| 70HRT15-count-cluster-clusterAnalysis.csv : 70HRT14-count-cluster-clusterAnalysis.csv | AACCGCAAGCAACACCCAGCAAGAAACATCCGACGCACGACGGGAGAAAGTGCATTACCACGATGTCGAT | 0.644 |
| 70HRT15-count-cluster-clusterAnalysis.csv : 70HRT14-count-cluster-clusterAnalysis.csv | AAGTCGTGTGATTGAATTGTCCCGACAGGTATCCGAAAGGGCACGGGATAAGCTCTGTATGGAGTCGAAT | 2.698 |
| 70HRT15-count-cluster-clusterAnalysis.csv : 70HRT14-count-cluster-clusterAnalysis.csv | ACCAAATCCCGAACTACAAATCCGAACGCTAACGGGACAATTGCGAAATGGAACATACGGGCCTGGTTGA | 1.788 |
| 70HRT15-count-cluster-clusterAnalysis.csv : 70HRT14-count-cluster-clusterAnalysis.csv | ACCAAGATAAACCGAGGTGTAAACGGGAAAACACATGGAATGACGTGCTGTAGTAGGTGTGATCACTGC | 110.032 |

Figure 39: FASTAptameR-Cluster_Enrich Output.

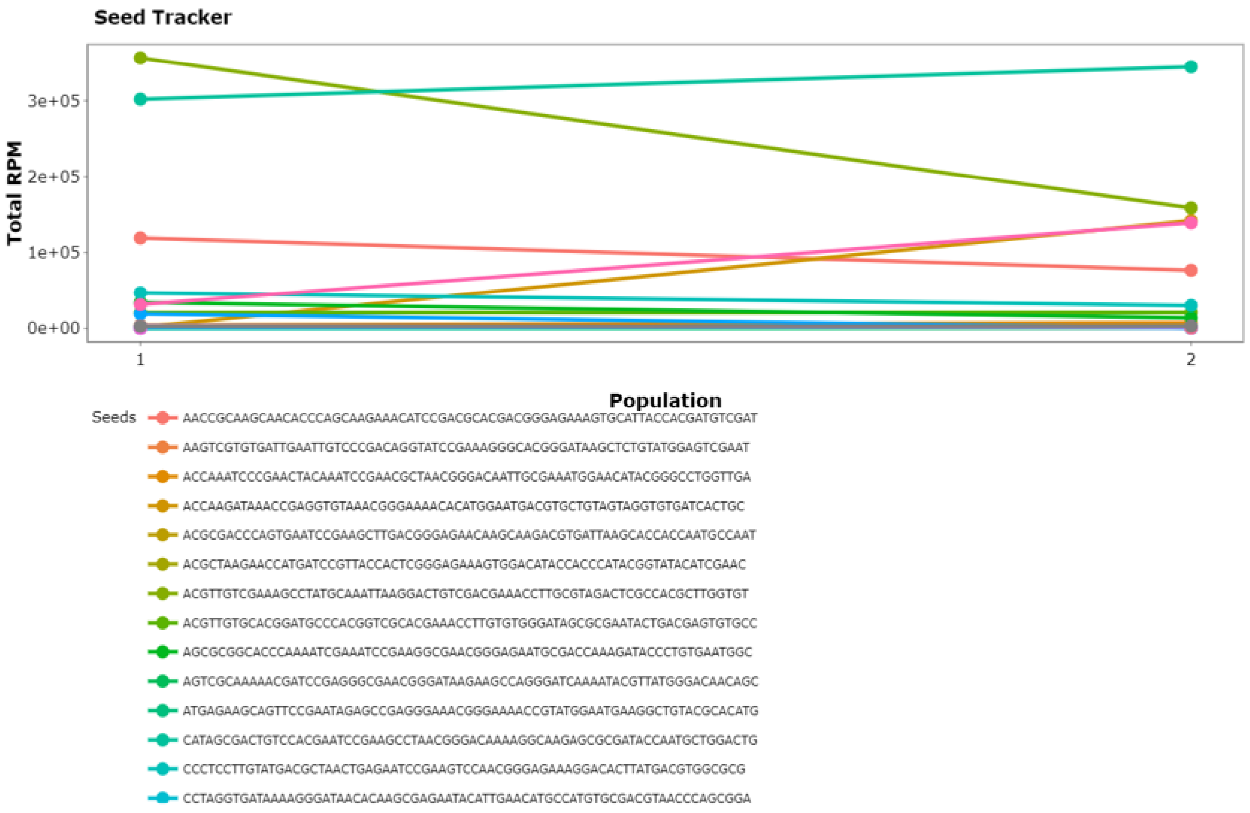

Figure 40: Cluster Seed Tracker Line Plot.

### 4 Version history
